# An auxin homeostat allows plant cells to establish and control defined transmembrane auxin gradients

**DOI:** 10.1101/2024.02.07.579341

**Authors:** Markus Geisler, Ingo Dreyer

## Abstract

Extracellular auxin maxima and minima are important to control plant developmental programs. Auxin gradients are provided by the concerted action of proteins from the three major plasma membrane auxin transporter classes AUX1/LAX, PIN and ABCB transporters. But neither genetic nor biochemical nor modelling approaches have been able to reliably assign the individual roles and interplay of these transporter types.
Based on the thermodynamic properties of the transporters, we show here by mathematical modeling and computational simulations that the concerted action of different auxin transporter types allow the adjustment of specific transmembrane auxin gradients. The dynamic flexibility of the “auxin homeostats” comes at the cost of an energy-consuming “auxin cycling” across the membrane.
An unexpected finding was that functional ABCB-PIN coupling appears to allow an optimization of the trade-off between the speed of auxin gradient adjustment on the one hand and ATP consumption and disturbance of general anion homeostasis on the other.
In conclusion, our analyses provide fundamental insights into the thermodynamic constraints and flexibility of transmembrane auxin transport in plants.

**Plain language summary:** The phytohormone auxin controls essentially plant development. Plant cells produce auxin and export it to establish patterns by local auxin minima and maxima. Although several transporter proteins are known to contribute to this process, the mechanism by which a defined auxin gradient can be produced is not clear. This study now uses mathematical modeling based on the thermodynamic features of the auxin transporters to illustrate in computational simulations the fundamental characteristics of an “auxin homeostat”. The concerted interplay of different auxin transporters allows plant cells to establish defined transmembrane auxin gradients that are the indispensable basis for polarized auxin maxima and minima and auxin fluxes within tissues.

## Introduction

Auxin is generally considered as a plant hormone that regulates various – if not all – aspects of plant growth and development, including cell elongation and expansion, tropisms, organogenesis and developmental patterning, vascular tissue differentiation, apical dominance and wound responses (for reviews, see Vanneste & Friml, 2009; Friml, 2021; Hammes *et al*., 2022). In the last decades, several independent approaches provided good evidence that the generation of extracellular (or apoplastic) auxin maxima (Benková *et al*., 2003; Robert & Friml, 2009) (and minima Sorefan *et al*., 2009) are primarily instructive for essential switches controlling these physiological and developmental programs, suggesting in principle a more morphogen-like mode of action (Geisler, 2021).

The generation of these apoplastic auxin maxima (and minima) involves a complex interplay of various processes, including local auxin biosynthesis and the cell-to-cell movement of auxin, called polar auxin transport (PAT). The latter is provided by the concerted action of specific transport proteins residing in the plasma membrane (PM), which were originally postulated in the 1970s in the so-called chemiosmotic hypothesis of auxin transport (Rubery & Sheldrake, 1973, 1974; Katekar & Geissler, 1977). It was based on the chemical nature of IAA (the major relevant auxin), which as a weak acid (pKa = 4.85) can partially cross the PM from the apoplast (pH 5.5) but not from the neutral (pH 7) cytoplasm requiring an export system (Zazímalová *et al*., 2010). It was thus postulated that auxin is transported into and out of the cell through the action of specific transport proteins in the PM (Rubery & Sheldrake, 1973, 1974; Raven, 1975; Goldsmith, 1977).

A number of auxin-transporting proteins have since been identified at the molecular level that belong to several evolutionary distinct families, with the AUXIN-RESISTANT1 (AUX1)/LIKE AUX1 (AUX1/LAX), PIN-FORMED (PIN) and ATP-BINDING CASSETTE subfamily B (ABCB) proteins being the best described (for latest reviews, see Hammes *et al*., 2022; Hammes & Pedersen, 2024). (i) AUX1/LAX are secondary active IAA/H^+^ symporters (Marchant *et al*., 1999; Kramer, 2004; Rutschow *et al*., 2014) that depend on the *proton motif force* (*pmf*) generated by the H^+^-ATPases and might be either electroneutral or electrogenic depending on the proposed H^+^/IAA^-^ ratios of 1 or 2. Recently, based on experimental and *in silico* data it was estimated that AUX1 is responsible for 75% of the PM influx, while 20% and 5% were attributed to other transporters and diffusion, respectively (Rutschow *et al*., 2014). (ii) So-called long/PM-based PINs are IAA^-^ exporters that are thought to be strictly dependent on the electrical component of the electrochemical gradient created by the H^+^-ATPases. However, despite recent publications of three valid PIN structures (Ung *et al*., 2022; Su *et al*., 2022; Yang *et al*., 2022; Hammes & Pedersen, 2024), the exact mechanism and energization of transport via PINs remains to be clarified. If IAA^-^ would act as a substrate, PINs can be considered as electrogenic uniporters (TC 2.A.1.), which is the most likely case, while the electroneutral substrate IAAH would make them facilitators (TC 1.A.8.). Based on the predominant, polar expression of some members, PINs are currently seen as the polar component contributing to polar auxin transport (Zazímalová *et al*., 2010; Geisler, 2021; Hammes *et al*., 2022; Hammes & Pedersen, 2024). (iii) ABCBs (TC 3.A.1.) are primary active (ATP-dependent) transporters that in most cases pump IAA^-^ out of the cell (Geisler *et al*., 2005; Geisler & Murphy, 2006; Zhang *et al*., 2018; Hao *et al*., 2020). However, two isoforms (ABCB4 and 21) were described as facultative im-/exporters (Santelia *et al*., 2005; Terasaka *et al*., 2005; Kamimoto *et al*., 2012). Importantly, due to their primary active energization, auxin translocation by ABCBs is independent of the electrochemical gradient and can thus work even against a very steep auxin gradient (Geisler & Murphy, 2006).

In direct dependence of their transport mechanism and energization, members of these three major families differentiate in their turn-over numbers and substrate specificity. While AUX1/LAX and PINs can be considered as relatively fast with estimated turnover numbers of 100-1.000 IAA/s, ABCBs that hydrolyze 2 ATP molecules per reaction cycle are slower (1-100 IAA/s). Overlapping expression profiles between major PIN and ABCB isoforms (Mellor *et al*., 2022) suggested a coordinated action between these transporter types. Co-immunoprecipitation and co-expression analyses using radiolabeled tracers and electrophysiological studies suggested that ABCB-PIN pairing increased transport capacity, resulting in synergistic transport rates that are greater than the sum of the transport rates of the individual transporters (Bandyopadhyay *et al*., 2007; Blakeslee *et al*., 2007). Interestingly, such cooperative, interactive ABCB-PIN coupling was found to also enhance substrate specificity (Blakeslee *et al*., 2007; Spalding, 2013; Geisler *et al*., 2017; Geisler, 2018). However, a genetic and biochemical dissection of the individual roles of ABCB and PIN classes by means of co-expression in heterologous systems resulted in unclear or even conflicting results, inspiring *in silico* investigations of these auxin transporters and their interplay at the root-tissue level. PIN-, ABCB- and PIN/ABCB-mediated auxin fluxes were modelled based on experimentally determined PIN and ABCB localization (Kramer, 2008; Band *et al*., 2014; Mellor *et al*., 2022). Although the results indicated that ABCBs function independently of PINs, only regulatory, specific ABCB-PIN interactions can reproduce experimentally observed auxin distributions (Mellor *et al*., 2022).

At present, we are left with the surprising finding that in the model plant *Arabidopsis thaliana* auxin gradients across the PM are apparently generated by a mixture of 20 primary and secondary active transporters (Geisler, 2021). However, neither modelling (Kramer, 2008; Band *et al*., 2014; Mellor *et al*., 2022) nor genetic and biochemical data (Blakeslee *et al*., 2007; Deslauriers & Spalding, 2021) allowed us to reliably assign the individual roles and interplay of all three major PM auxin transporter classes in the establishment of transmembrane auxin gradients. Related to this uncertainty is the open, puzzling question of why cells appear to employ two classes of auxin exporters, including ABCBs, which pump IAA^-^ out of the cell along its gradient at the expense of ATP consumption.

In order to address these pressing questions from a different angel, we here present an *in-silico* approach. In contrast to previous computational studies aiming at modeling auxin-mediated plant development (Prusinkiewicz, 2004), signaling (Rutten *et al*., 2022) or mid-range transport (Band et al. 2014, Mellor et al. 2022), we have focused on auxin membrane transport and have chosen an approach based on the thermodynamic properties of the transporters. This type of approach has recently been successfully implemented to identify and characterize K^+^, anion and Ca^2+^ homeostats in the plasma and vacuolar membranes of plant cells (Dreyer, 2021; Dindas *et al*., 2021; Dreyer *et al*., 2022; Li *et al*., 2024). Homeostats are networks of different types of transporters that are permeable for a specific ion species, either alone or together with other ions, e.g., H^+^. The fluxes via these transporters are coupled to each other via the membrane voltage and the respective transmembrane gradients of the permeating ion(s). As a consequence, homeostats have new dynamic properties that enable the control of ion fluxes across a membrane in a concerted action, allowing for example the regulation of ion homeostasis and osmoregulation. The gain of flexibility in adjusting transmembrane concentration gradients is inevitably accompanied by the establishment of energy-consuming transport cycles at the membrane (Dreyer, 2021).

Our results on auxin transporter networks now suggest that a combination of two different transport processes for auxin, e.g. diffusion and export, already forms an efficient auxin homeostat. This means that the cell could adjust a specific transmembrane auxin gradient by adapting transporter activities. Further elaborated homeostats with different mixtures of AUX1/LAX, PIN, and ABCB transporters exhibited particular fine-tuned properties. For example, the use of two different transporter types for IAA^-^ export, PIN and ABCB, mitigated the drawbacks of the individual transporters in terms of specificity and speed, respectively. The analyses presented here provide fundamental insight into the thermodynamic flexibility of transmembrane auxin transport in plants.

## Materials and Methods

### Mathematical Description of Transport Processes

#### Proton pump, potassium homeostat and anion homeostat

The mathematical description of the H^+^-ATPase and the transporters involved in the K- and the A-homeostat is detailed in the pilot study on homeostats modeling (Dreyer, 2021). For completeness, the equations are summarized in the following:

H^+^-ATPase:

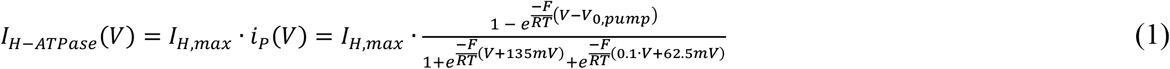

K_+_ channel (flux *J*_*K,KC*_, current *I*_*KC*_):

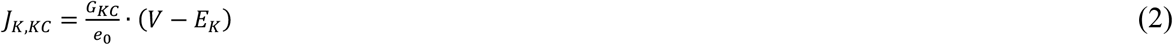

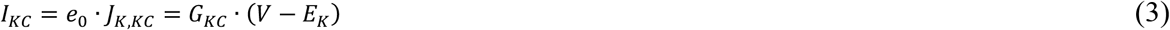

1H_+_:1K_+_ symporter (fluxes *J*_*H,KHs*,_ *J*_*K,KHs*,_ current *I*_*KHs*_):

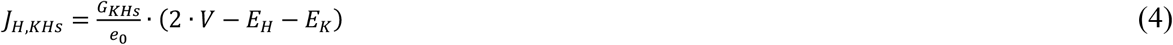

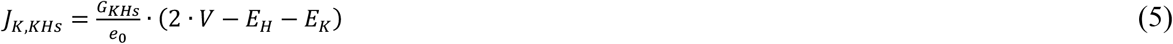

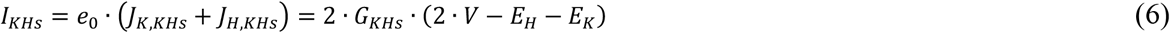

1H_+_:1K_+_ antiporter (fluxes *J*_*H,KHa*,_ *J*_*K,KHa*_):

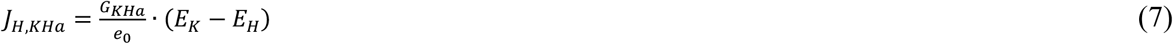

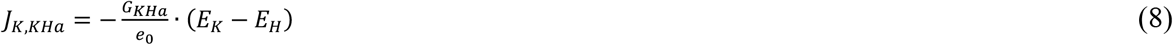

anion channel (flux *J*_*A,AC*_, current *I*_*A*_):

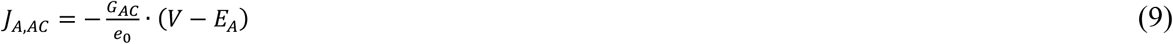

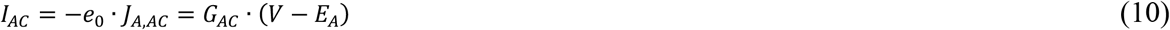

2H_+_:1 anion symporter (fluxes *J*_*H,HA*,_ *J*_*A,HA*,_ current *I*_*HA*_):

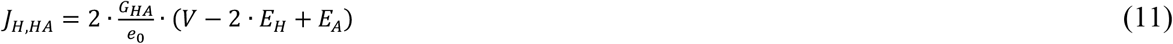

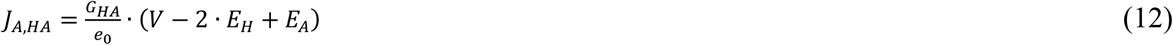

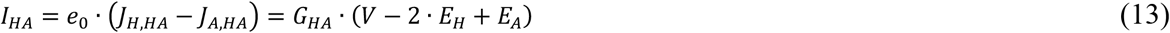

where *V* is the membrane voltage, *e*_*0*_ the elementary charge, and *E*_*K*_, *E*_*A*_, and *E*_*H*_ the Nernst voltages for potassium, the anion (Cl^-^ or/and NO_3-_) and for protons, respectively. *G*_*KC*_, *G*_*KHs*_, *G*_*KHa*_, *G*_*AC*_, *G*_*HA*_ are the membrane conductance of the respective transporter type (unit pA/V). *I*_*H,max*_ (unit pA) is the maximum H^+^-pump current of the cell, while *V*_*0,pump*_ is the voltage at which the pump current is zero. In the examples presented in this study, this value was set exemplarily to *V*_*0,pump*_ = -200 mV; different values would not qualitatively change the results. *F* is the Faraday constant, *R* the gas constant and *T* the absolute temperature.

#### Protonation of auxin and passive diffusion across the membrane

Depending on the prevalent pH, the auxin indole-3-acetic acid exists as anionic base IAA^-^ or its conjugate acid, IAAH. The reaction [*IAA*^−^] + [*H*^+^] ⇌ [*IAAH*] has a pKa of 4.75. While the lipid bilayer can be considered as impermeable for the negatively charged form, the uncharged protonated form is able to diffuse across the membrane with a net efflux of:

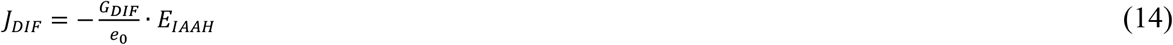

with 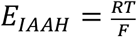· ln 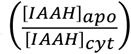 and the apoplastic and cytolic IAAH concentrations [IAAH]_apo_ and [IAAH]_cyt_. Considering the (de)protonation of auxin, *E*_*IAAH*_ can be expressed as *E*_*IAAH*_ = *E*_*H*_ -*E*_*IAA*_, with 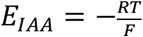· ln 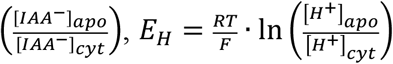 and the apoplastic and cytosolic IAA^-^ and H^+^ concentrations [IAA^-^]_apo_, [IAA^-^]_cyt_, [H^+^]_apo_, and [H^+^]_cyt_. Although the electroneutral diffusion is not connected with electrical current, an electrical conductance, *G*_*DIF*_ (pA/V) can be defined. This trick has technical reasons to unify the equations.

1 IAA^-^ : 1 H^+^ symporter, AUX

Proton (*J*_*H,AUX*_) and IAA^-^ (*J*_*IAA,AUX*_) net flux from the cell via a 1:1 IAA^-^/H^+^ symporter:

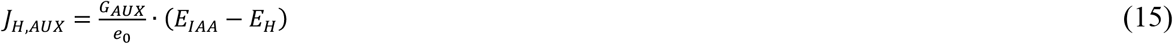

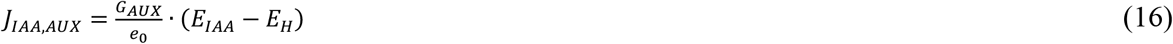

Also here, an electrical conductance, *G*_*AUX*_ (pA/V), was defined analogously to the other transport processes.

IAA^-^ uniporter, PIN

The mechanism of direct uniport of IAA^-^ anions across membranes by PIN transporters has not yet been clarified. The IAA^-^ flux (*J*_*IAA,PIN*_) might be driven only by the electrochemical IAA^-^ gradient or provoked by flux coupling with an anion flux (*J*_*A,PIN*_) driven by a mixed electrochemical driving force *E*_*mixed*_ = β·*E*_*IAA*_ + (1-β)·*E*_*A*_. To cover all possibilities, fluxes and currents were described as:

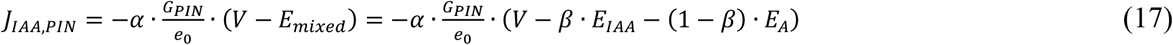

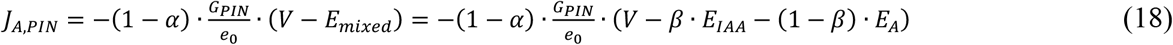

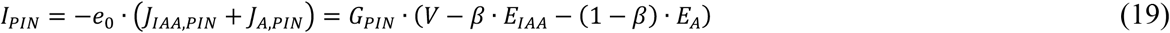

Here, *G*_*PIN*_ (pA/V) represents the conductance of the auxin-permeable channel, 0 ≤ α ≤ 1 denotes the fraction of the auxin flux in the total flux, i.e. the real permeability for IAA^-^, and 0 ≤ β ≤ 1 indicates in as much IAA^-^ influences the zero-current voltage of the transporter, which is widely used to calculate its “relative permeability” (Navarro-Retamal *et al*., 2021). The parameter pair α = 1 and β = 1 represents an auxin-selective PIN, while with α = 0 it would not be permeable for auxin.

#### ABCB transporter

An ATP binding cassette transporter of subfamily B (ABCB) that pumps IAA^-^ from the cytosol to the apoplast was described similar to an H^+^-ATPase (Dreyer, 2017; Reyer *et al*., 2020) by a multiple-state model, resulting in approximated mathematical expressions for the current-voltage relationship and the flux:

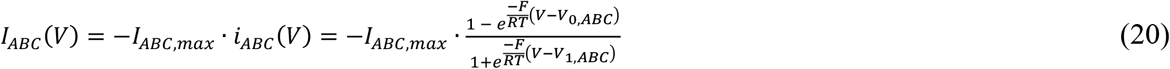

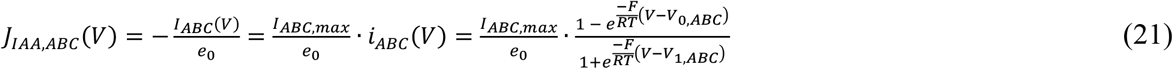

Here, *I*_*ABC,max*_ (pA) is the maximum current, which depends on the number of active ABC transporters in the membrane and the cytosolic and apoplastic IAA^-^ concentrations. *V*_*0,ABC*_ is the voltage at which the transporter current is zero. It depends on the cytosolic and apoplastic proton concentrations and on the cytosolic ATP, ADP, and Pi concentrations, *i*.*e*., on the energy status of the cell. Because ABCB transporters consume 2 ATP molecules per transported substrate ion/molecule, this value is rather negative. In the examples presented in this study, this value was set exemplarily to *V*_*0,ABC*_ = -400 mV; different values would not qualitatively change the results. The parameter *V*_*1,ABC*_ > *V*_*0,ABC*_ indicates the midpoint of the S-type shaped curve of *i*_*ABC*_(*V*) (**Figure S1**). For the simulations in this study, we used *V*_*1,ABC*_ = -250 mV. Different values do not change the results qualitatively. A positive value of *I*_*ABC,max*_ describes an export of IAA^-^ (negative current, positive flux), while *I*_*ABC,max*_ < 0 represents an import (positive current, negative flux).

### Changes in voltage and concentrations and steady state conditions

The membrane voltage changes according to

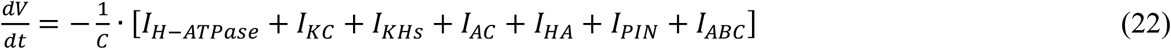

the cytosolic potassium concentration according to

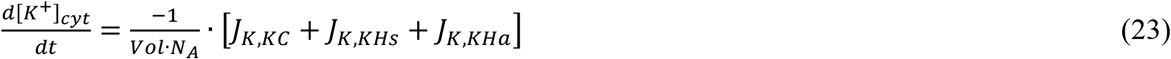

the cytosolic anion concentration (apart from IAA^-^) according to

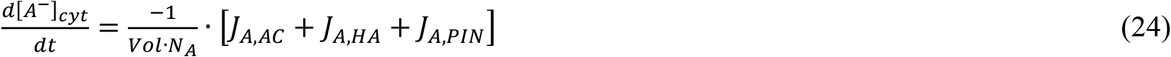

and the cytosolic auxin (IAA^-^) concentration according to

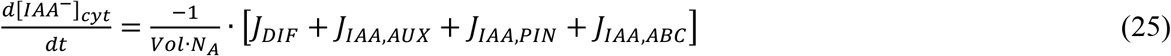

with the membrane capacitance *C*, the cellular volume *Vol*, and the Avogadro constant *N*_*A*_. The steady state analyzed in this study is defined by the conditions:

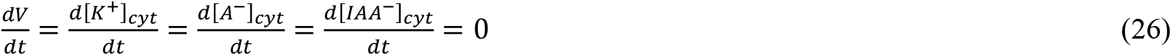

### Proton concentrations

Proton concentrations are predominantly determined by metabolic processes (Sanders & Slayman, 1982; Wegner & Shabala, 2020; Wegner *et al*., 2021). Therefore, *E*_*H*_ was fixed to +97.9 mV for the steady state analyses in this study. This value represented a pH gradient of pH_cyt_ = 7.2 and pH_apo_ = 5.5 and was chosen without any bias. A different ΔpH would qualitatively produce the same results. In steady-state, the net proton flux across the membrane is zero:

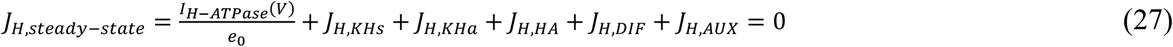

This result is implicitly included in the equations (22-26).

### Auxin gradient

In the figures, we present the transmembrane auxin gradient as the fraction 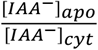, which is proportional to the total auxin gradient 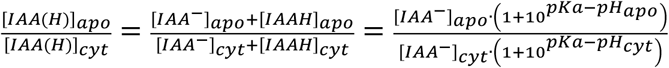. In our simulations (pH_cyt_ = 7.2, pH_apo_ = 5.5), the proportional factor is 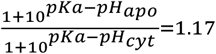 .

### Renormalization to eliminate a redundant parameter

Equations (22-25) were multiplied by the factor 1 = *I*_*H,max*_/*I*_*H,max*_. *I*_*H,max*_ was combined with the pre-factors to *I*_*H,max*_/*C* and *I*_*H,max*_/(*Vol*·*N*_*A*_), while 1/*I*_*H,max*_ was used to normalize the parameters to the maximal pump current: *i*_*P*_(*V*) = *I*_*H-ATPase*_(*V*)/*I*_*H,max*_ (without unit), *g*_*X*_ = *G*_*X*_ /*I*_*H,max*_ (X=KC, KHs, KHa, AC, HA, DIF, AUX, PIN; all *g*_*X*_ with unit V^-1^), *g*_*ABC*_ = *I*_*ABC,max*_/*I*_*H,max*_ (without unit). This renormalization eliminated one redundant parameter and yielded the final equation system for the steady state conditions (**Supplementary Material**).

### Numerical determination of steady state conditions

For each parameter set *g*_*x*_ the equation system has a steady state solution for the membrane voltage *V* and the *E*_*X*_-values. These were determined first by solving **Equation S1** and finally by numerically resolving the resultant implicit equations.

### Model assessment

It should be noted that all conclusions drawn in this study are independent of the exact description of the fluxes *J*_*X*_ (for further justification see Dreyer, 2021). The screening of the entire reasonable parameter space [0,∞) for the *g*_*X*_ guaranteed the coverage of all possible cases that a cell can achieve in the considered scenarios, and allowed us to assess the limits of the flexibility of the system under consideration in steady state, even without prior knowledge of specific transporter features.

## Results

To explore the existence of an auxin homeostat and to understand the contributions of different auxin transporters to auxin homeostasis, we adopted a stepwise computational approach. In simulating the involved transporter networks, we started with the simplest system and increased successively its complexity. The main goal was to determine the dependence of the steady state condition of the system (homeostasis) on the activity of the different transporters contributing to auxin transport. For this purpose, the proton concentrations could be kept constant and were set to physiological pH_cyt_ = 7.2 and pH_apo_ = 5.5. Other values would not change the presented results qualitatively.

### Reference condition

We investigated auxin transport in a physiological context of a cell with active H^+^-pumps, K^+^ and anion (A^-^, i.e. Cl^-^ and/or NO_3-_) transporters. As a reference condition, we chose settings for the K- and A-homeostats (Dreyer, 2021) that resulted in steady-state values of *V* = -150mV, *E*_*K*_ = - 120mV, *E*_*A*_ = 40mV and *E*_*H*_ = 98mV in the absence of auxin (for details see Supplementary Material). The adjusted parameter set of the K- and A-homeostats was then held constant when we tested the effect of the different auxin transporters on *V, E*_*K*_, and *E*_*A*_, and cytosolic and apoplastic auxin concentrations. An auxin transport-induced change in *E*_*K*_ and *E*_*A*_ (under constant external conditions) means a change in the internal K^+^ and anion concentration, respectively, which could have detrimental effects on other cellular processes. To illustrate these adverse effects, we defined exemplarily a 5%-change in internal concentrations caused by auxin transporters as a critical level. This is equivalent to changes of |Δ*E*_*K*_| ≥ 1.25 mV or |Δ*E*_*A*_| ≥ 1.25 mV.

#### Case 1: Passive diffusion across the membrane

In a first *‘what-if’* scenario (**Figure 1a**) we considered the simplest case, in which auxin can diffuse across the membrane in its protonated form (IAAH), while it gets trapped in the respective compartment in its anionic form (IAA^-^) (Zazímalová *et al*., 2010). Irrespective of the starting conditions, the diffusion process reached steady state when [IAAH]_cyt_ = [IAAH]_apo_ (**Figure 1c, d**, left, black curves). This does not mean, however, that also the IAA^-^ concentrations were the same on both sides of the membrane. Instead, the pH-dependent (de)protonation process resulted in 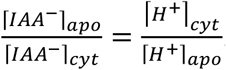. For a cytosolic pH of 7.2 and an external pH of 5.5 this implied [IAA^-^]_cyt_ = 50×[IAA^-^]_apo_ in steady state (**Figure 1c, d**, right, black curves). Different gradients could only be established by varying the cytosolic and/or extracellular pH conditions. Thus, auxin diffusion alone was not sufficient to effectively control the apoplastic auxin concentration (**Figure 1c, d**, black curves). Also, the electroneutral diffusion of auxin did not influence other transport processes. In particular, it has not changed *E*_*K*_ or *E*_*A*_.

**Figure 1.**
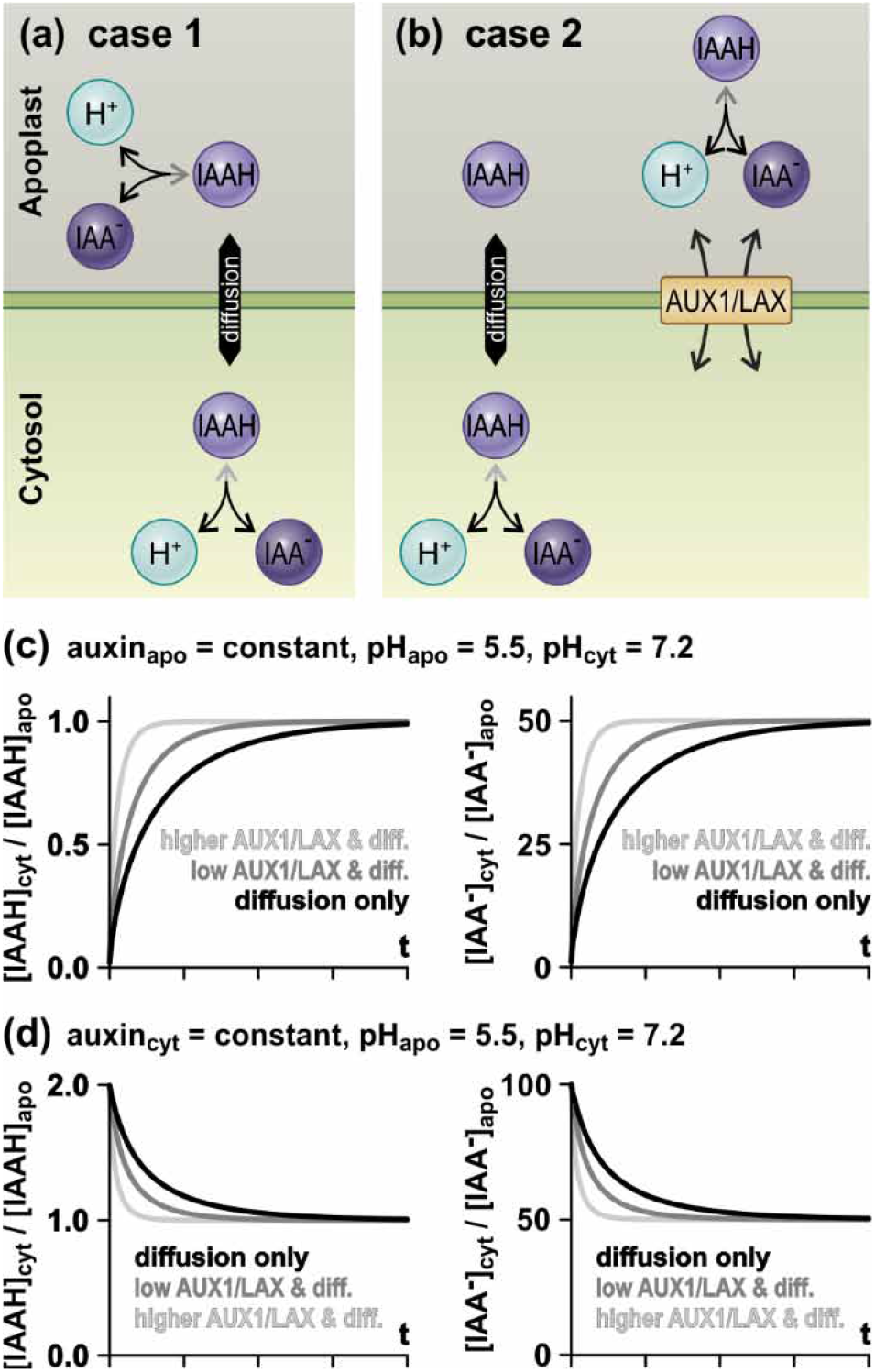
Diffusion and electroneutral transport of auxin are not sufficient to effectively control the apoplastic auxin concentration. (a) Case 1: The protonated form of auxin (IAAH) can passively diffuse across the membrane. (b) Case 2: In addition to diffusion, an IAA/H symporter (AUX1/LAX) enables electroneutral transmembrane transport of auxin. (c and d) Simulations of the systems shown in A (black curves) and B (purple curves) with pH_apo_ = 5.5 and pH_cyt_ = 7.2. (c) Equilibration starting from [IAAH]_cyt_ < [IAAH]_apo_ keeping the apoplastic auxin concentration constant. (d) Equilibration starting from [IAAH]_cyt_ > [IAAH]_apo_ keeping the cytosolic auxin concentration constant. The black curves (diffusion) represent the system exclusively with diffusion. For the dark grey curves (low AUX1/LAX activity) the diffusion process was supported by an electroneutral IAA^-^/H^+^ transport of the same magnitude as the diffusion, while for the faint grey curves (higher AUX1/LAX activity) the electroneutral IAA^-^/H^+^ transport was five times larger. Neither the diffusion nor AUX1/LAX affected the K- and A-homeostats in the cell (not shown).

#### Case 2: Diffusion and electroneutral IAA^-^/H^+^ symporter (AUX1/LAX)

In the next *‘what-if’* scenario we added an electroneutral IAA^-^/H^+^ 1:1 co-transporter (AUX1/LAX) to our model system (**Figure 1b**; for non-electroneutral ratios, see Discussion). This additional transport pathway allowed to modulate the speed of the diffusion process but did not change the steady state characteristics (**Figure 1c, d**, grey curves). The larger the AUX1/LAX activity was, the faster the steady was reached. Also here, the final gradients were [IAAH]_cyt_ = [IAAH]_apo_ and [IAA^-^]_cyt_ = 50×[IAA^-^]_apo_ as already observed in case 1 and depended only on the transmembrane pH-gradient. Thus, neither auxin diffusion alone nor in combination with an electroneutral AUX1/LAX was sufficient to effectively control the apoplastic auxin concentration. These transport processes also had no effect on the K- and A-homeostats, leaving *E*_*K*_ and *E*_*A*_ unaffected.

#### Cases 3 and 4: Auxin-transporting ABCB-transporter (ABCB)

In the next step, we extended the former scenario to include an auxin-transporting ABCB-transporter (**Figure S1**) that pumps IAA^-^ either along its gradient out of the cell (case 3; **Figure 2a**) or — as shown in some conditions (Kamimoto *et al*., 2012) — against its electrochemical gradient into the cell (case 4; **Figure S2a**). In both cases, the hydrolysis of ATP provided the energy for substrate transport. Remarkably, this additional transporter decoupled the dependence of the transmembrane auxin gradient on the prevailing pH conditions. To explain this for case 3, we start in Figure 2 from a condition, in which the activity of the ABCB transporter was zero, resulting in an auxin gradient of [IAA^-^]_cyt_/[IAA^-^]_apo_ = 50 (*g*_*ABC*_ = 0, **Figure 2b**, magenta curve) as in cases 1 and 2. Now, when the activity of the exporting ABCB-transporter increased, the steady-state value of the auxin gradient decreased with elevating activity levels (*g*_*ABC*_ > 0). The new steady-state values depended not only on the activity of the ABCB-transporter (*g*_*ABC*_) but also on that of the diffusion process (*g*_*DIF*_+*g*_*AUX*_). Thus, due to the combination of the two different transport processes, diffusion and primary active transport, the cell gained control over the transmembrane auxin gradient. The auxin gradient could now be adjusted by tuning the activity of the involved transporters (AUX1/LAX and ABCB). Nevertheless, this advantage came at the inevitable expense of energy-consuming “auxin cycling” (transporter-mediated im- and export) across the membrane (**Figure 2a**) as already shown for K^+^ and anion homeostats (Dreyer, 2021). The permanent efflux of IAA^-^ via the ABCB-transporter was compensated in steady-state by an IAAH/H^+^-IAA^-^ influx by diffusion or via AUX1/LAX (**Figure 2a**).

**Figure 2.**
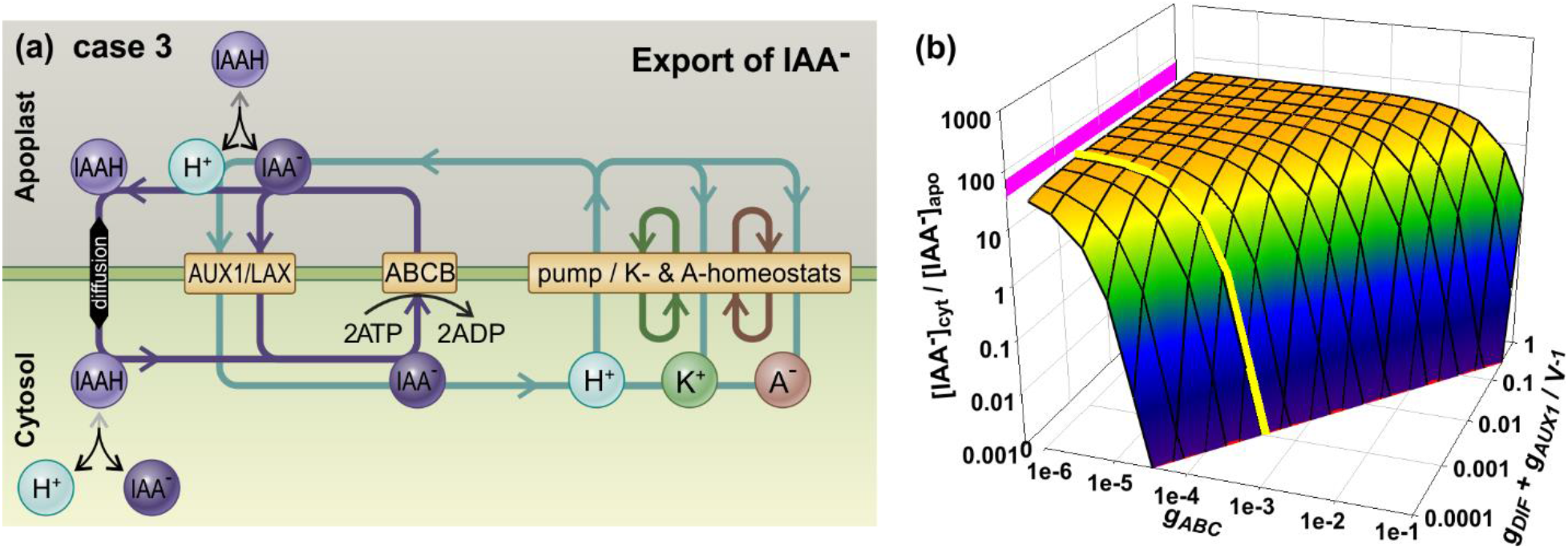
IAA^-^ exporting ABCB-transporters can decrease the transmembrane auxin gradient (case 3). (a) Schematic representation of the transmembrane transport processes. ABCB-transporters export IAA^-^, which is compensated in steady state by IAAH/H^+^-IAA^-^ influx by diffusion or via AUX1/LAX. The involved proton fluxes in this cycling feeds back on the H^+^-pump dependent homeostats for potassium (K^+^) and anions (A^-^). Substrate cycling is indicated by the colored arrows. (b) Transmembrane [IAA^-^]_cyt_/[IAA^-^]_apo_ gradient in steady state. The magenta line shows the state in the absence of ABCB transporter activity, which is determined by diffusion of IAAH and pH-dependent (de)protonation reactions. The yellow line depicts values for which *g*_*DIF*_*+g*_*AUX1*_ = 10^-3^ V^-1^ to illustrate exemplarily a lower limit determined by transporter-independent IAAH-diffusion across the membrane (*g*_*AUX*_ = 0 V^-1^).

In contrast to an exporting ABCB transporter (case 3) that was able to overcompensate for the diffusion-induced reflux of IAAH into the cell, an importing transporter was able to increase cytosolic auxin beyond the diffusion limit (case 4; **Figure S2**). Also here, the auxin gradient was [IAA^-^]_cyt_/[IAA^-^]_apo_ = 50 in the absence of ABCB-transporter activity (*g*_*ABC*_ = 0, **Figure S2b**, magenta curve). However, the steady-state value of the auxin gradient raised with elevating activity levels (|*g*_*ABC*_| > 0). In this case, the permanent influx of IAA^-^ via the importing ABCB-transporter was compensated in steady-state by an IAAH/H^+^-IAA^-^ efflux by diffusion or via AUX1/LAX (**Figure S2a**).

In both cases 3 and 4, the involved proton fluxes affected the other homeostats in the membrane. The larger was the ATP-driven IAA^-^ export in case 3, the larger was the H^+^ influx mediated by IAAH diffusion and/or AUX1/LAX-mediated H^+^/IAA^-^ symport. This proton influx partially dissipated the *pmf* established by H^+^-ATPases. As a consequence, the membrane voltage became less negative (**Figure S3a**), cytosolic K^+^ decreased (E_K_ less negative, **Figure S3b**) and cytosolic anion concentration increased slightly (E_A_ more positive, **Figure S3c**). In case 4 (**Figure S2**), the greater was the ATP-driven IAA^-^ import, the greater was the H^+^ efflux mediated by AUX1/LAX/diffusion. This, in turn, caused a more negative membrane voltage in the model scenario (**Figure S3d**), higher cytosolic K^+^ (E_K_ more negative, **Figure S3e**) and a slightly lower cytosolic anion concentration (E_A_ less positive, **Figure S3f**). Thus, in case 4, the energy consumed by the IAA^-^-pumping ABCB-transporter was partially converted into an additional proton-motive force (*pmf*) that exerted its effect on K- and A-homeostasis. In summary, the employment of ABCB-transporters enabled the control of the transmembrane auxin gradient but the process of homeostatic control consumed energy and affected K^+^ and anion homeostasis.

#### Cases 5 and 6: Auxin uniporter (PINs)

An alternative to the active export of auxin via ABCB-transporters could be the passive efflux of IAA^-^ along its electrochemical gradient. To evaluate this possibility, we simulated a new scenario and replaced the ABCB-transporter in cases 3-4 with an auxin-permeable uniporter (PIN, **Figure 3a**). The mechanism of direct uniport of IAA^-^ anions across membranes by PIN transporters is not yet clear, despite recent structural insights (Hammes *et al*., 2022; Ung *et al*., 2022; Hammes & Pedersen, 2024). Therefore, we chose a broad-range approach that covered the entire spectrum between IAA^-^-selective (case 5) and non-selective, auxin-permeable PIN anion channels (case 6). To do so we introduced two parameters: (i) α (0 ≤ α ≤1) described the real permeability of PIN for IAA^-^ in relation to other anions such as Cl^-^ or NO_3_^-^, and (ii) β (0 ≤ β ≤1) described the influence of IAA^-^ on the reversal voltage of PIN, i.e. the driving force for transport. An IAA^-^-selective PIN (case 5) was characterized by (α = 1) while α < 1 represented a PIN that allowed also the passage of other anions (case 6; for more details see Material and Methods). In each of these cases, like the ABCB-transporter, PIN also converted the static case 2 (**Figure 1**) into a dynamic scenario, in which the transmembrane auxin gradient could be adjusted by fine-tuning the activities of PIN and/or AUX1/LAX (case 5: **Figure S4**, case 6: **Figures 3b-e** and **S5**). In its consequences, PIN was very similar to the IAA^-^-exporting ABCB-transporter (case 3, **Figure 2**) and could not increase the auxin gradient beyond the diffusion-determined limit (cyan lines in **Figures 3b, S4a**,**e**,**i** and **S5a**,**e**,**i**).

**Figure 3.**
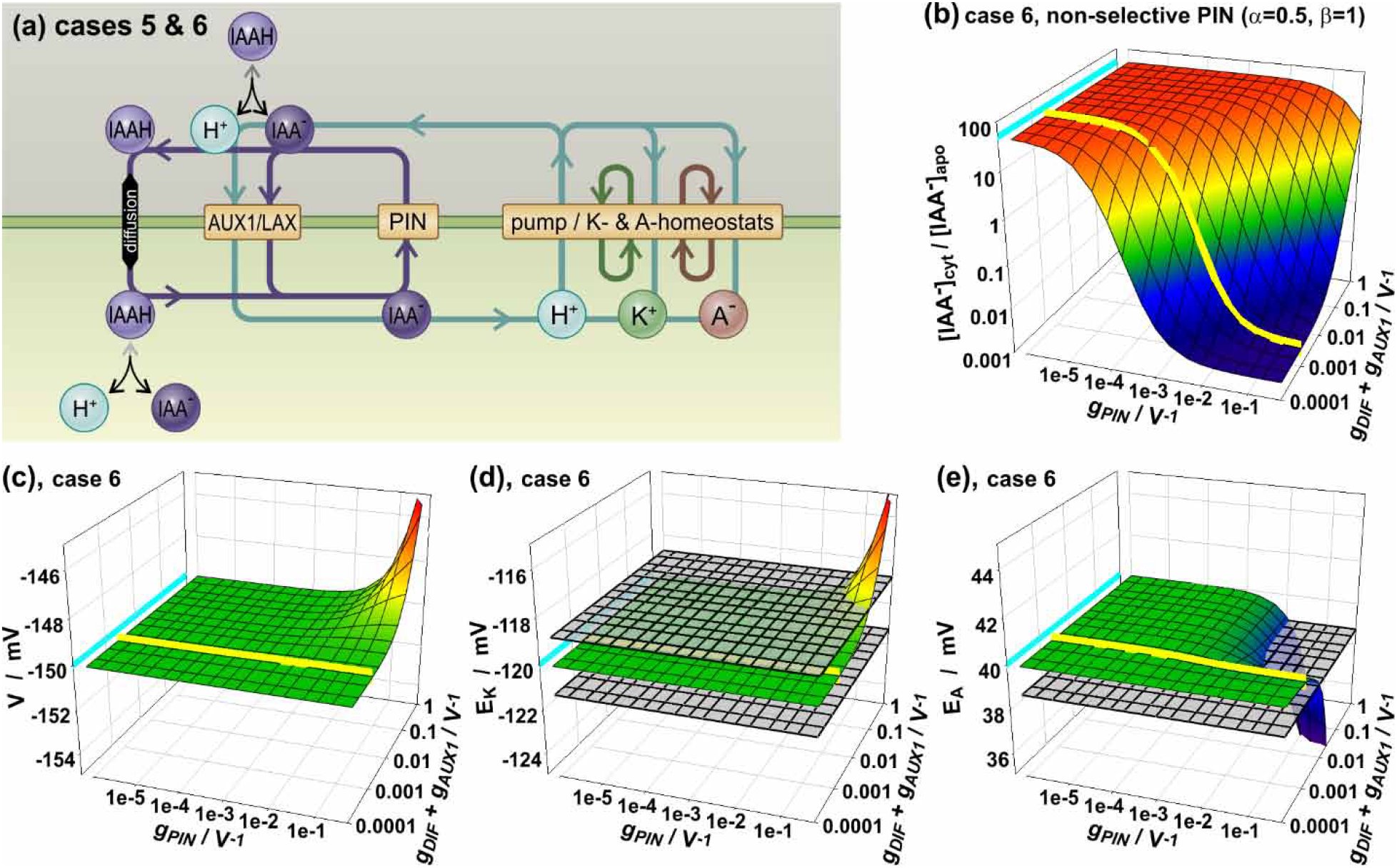
PIN transporters can control the transmembrane auxin gradient. (a) Schematic representation of the transporter network of cases 5 and 6, which include PIN and AUX1/LAX in addition to the natural IAAH diffusion and the K and A homeostats and the proton pump. The colored arrows and cycles indicate the fluxes via the different transport pathways in steady state. (b-e) Transmembrane [IAA^-^]_cyt_/[IAA^-^]_apo_ gradient (b), membrane voltage, *V* (c), and equilibrium voltages for K^+^, *E*_*K*_ (d), and anions, *E*_*A*_ (e) in steady state for a case 6 scenario (non-selective PIN, parameters α = 0.5 and β = 1) as a function of the PIN and AUX1/LAX/diffusion transporter activities. The cyan lines show the steady state in the absence of PIN transporter activity, which is determined by diffusion of IAAH and pH-dependent (de)protonation reactions. The yellow lines depict values for which *g*_*DIF*_*+g*_*AUX1*_ = 10^-3^ V^-1^ to illustrate exemplarily a lower limit determined by transporter-independent IAAH-diffusion across the membrane (*g*_*AUX*_ = 0 V^-1^).

Like in every homeostat, also in the scenarios 5 and 6 the control over the transmembrane auxin gradient was accompanied by a permanent, energy-consuming auxin cycling across the membrane. After steady state was established, any IAA^-^ released by PIN was retrieved either as IAAH by diffusion or as H^+^/IAA^-^ symport via AUX1/LAX. The involved proton influx dissipated part of the *pmf* established by the H^+^-ATPase (**Figure 3a**). As a consequence, high activity of PIN depolarized the membrane, which in turn affected K^+^ (E_K_) and anion (E_A_) homeostasis. Interestingly, we observed qualitative and quantitative differences in this respect that depended on the selectivity of the PINs (**Figures 3, S4, S5**). While the parameter β had only a minor effect on homeostasis (**Figures S4, S5**), the parameter α, i.e. the real permeability, was critical, in particular for anion homeostasis (**Figure S6**). To gain better insight into the effect of PIN selectivity on anion homeostasis, we compared case 4, in which an ABCB transporter exports IAA^-^, to case 5 with an IAA^-^-selective PIN (α = 1) and case 6 with a non-selective PIN (α < 1). We calculated the deviation of E_A_ for a determined [IAA^-^]_cyt_/[IAA^-^]_apo_ gradient at a given diffusion/AUX1/LAX activity (**Figure 4**). Both the ABCB transporter and the IAA^-^ selective PIN affected the anion gradient (E_A_) in the same moderate manner when adjusting the same transmembrane auxin gradient (**Figure 4a,b**). When a strong IAA^-^ efflux was required (*i*.*e*., low [IAA^-^]_cyt_/[IAA^-^]_apo_ ratios), the cytosolic anion concentration [A^-^]_cyt_ increased slightly, in particular at high IAAH/H^+^-IAA^-^ diffusion rates (i.e., larger *g*_*DIF*_+*g*_*AUX*_ values). The non-selective PIN, in contrast, interfered more severely with general anion homeostasis (**Figure 4c**). Now, [A^-^]_cyt_, and thus E_A_, decreased strongly even at moderate diffusion rates and also at higher [IAA^-^]_cyt_/[IAA^-^]_apo_ ratios. This detrimental effect was anion specific as the three compared cases did not differ remarkedly in the influence of the auxin homeostat on membrane voltage or the potassium gradient (E_K_; **Figure S7**). Thus, when PIN was not exclusively selective for auxin (α < 1, case 6), the activity of PIN had a detrimental effect on the steady state of the anion A^-^, E_A_ (compare **Figure S6d** with **S6h**,**l**,**p**).

**Figure 4.**
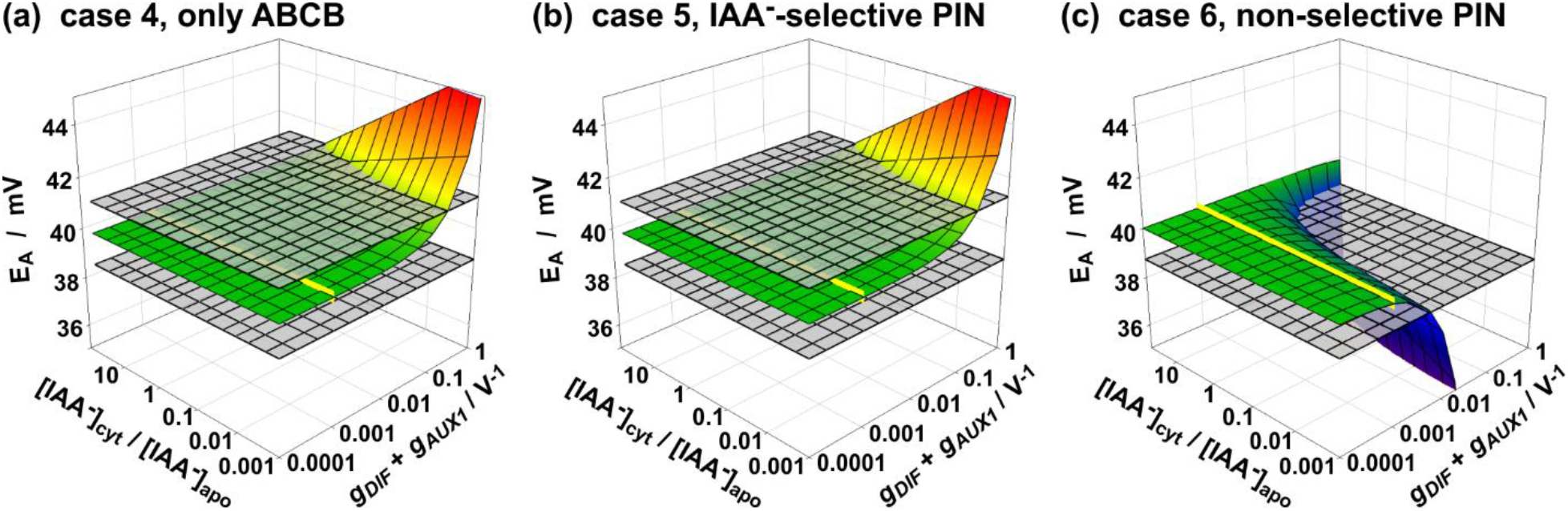
Non-selectivity of PIN can have detrimental effects on the transmembrane anion gradient of the cell. Comparison of the steady values of 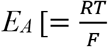 · ln 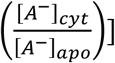 depending on auxin diffusion (*g*_*DIF*_+*g*_*AUX1*_) and the desired transmembrane auxin gradient ([IAA^-^]_cyt_/[IAA^-^]_apo_) in case 4 with IAA^-^-exporting ABCB transporters (a), case 5 (α = 1, β = 1) with IAA^-^-selective PINs (b), and case 6 (α = 0.5, β = 1) with non-selective PINs (c). The grey surfaces indicate thresholds at which the transmembrane anion gradient changes by 5% (|Δ*E*_*A*_| = 1.25 mV) compared to the reference condition without auxin transporters. The yellow line indicates values for which *g*_*DIF*_*+g*_*AUX1*_ = 10^-3^ V^-1^ to illustrate exemplarily a lower limit determined by transporter-independent IAAH-diffusion across the membrane (*g*_*AUX*_ = 0 V^-1^).

As suggested by an experimental study (Deslauriers & Spalding, 2021) and in view of known selectivity mechanisms (Gouaux & MacKinnon, 2005), e.g. for K^+^ channels (Doyle *et al*., 1998), it is rather reasonable to postulate that a channel or uniporter-like PIN has difficulties in excluding small anions such as Cl^-^ or NO_3_^-^ while transporting the relatively large IAA^-^. Usually, a high selectivity for larger substrates is achieved by conformational changes of the transporter, as in the case of ABC transporters (Srikant & Gaudet, 2019). Therefore, α < 1 (non-selective PIN) is more realistic than α = 1 (IAA^-^-selective PIN).

#### Case 7: A combination of ABCB and PIN can mitigate the drawbacks of the individual transporters

Exporting ABCB transporters and PINs function redundantly in their role of releasing IAA^-^ from the cytosol to the apoplast. Still, using one transporter or the other involves different tradeoffs for the cell. The primary active transport of IAA^-^ by the ABCB transporter is rather slow (estimated turnover numbers range between 1 – 100 IAA^-^/s) and needs the energy from ATP hydrolysis. But the conformational changes of the protein during the pumping process allow the larger IAA^-^ to be preferred over the smaller Cl^-^ and NO_3_^-^ anions. In contrast, the most likely uniport mechanism of PIN allows for rapid transport (100 - 1.000 IAA^-^/s), which inevitably comes at the expense of selectivity between IAA^-^ and smaller anions (Deslauriers & Spalding, 2021).

The similarities and differences of ABCB and PIN provoked the question whether the combination of both “imperfect” transporter types may result in beneficial effects. We therefore designed a new scenario that combined all transporters, AUX1/LAX, ABCB, and PIN (case 7; **Figure 5**). As in the cases 4 to 6, this transporter network allowed the transmembrane auxin gradient to be set by regulating the transporter activities. Again, the homeostatic control came at the expense of energy-consuming auxin cycling across the membrane, which also affected K^+^ and anion homeostasis (**Figure 5a**). However, these effects were not always equally pronounced. For the purpose of illustration, we fixed the diffusion/AUX1/LAX component (*g*_*DIF*_+*g*_*AUX1*_ = 0.05 V^-1^) and varied the activities of PIN, *g*_*PIN*_, and ABCB, *g*_*ABC*_. A desired auxin gradient (*i*.*e*., a chosen [IAA^-^]_cyt_/[IAA^-^]_apo_ ratio) could be achieved by many (*g*_*PIN*_|*g*_*ABC*_) pairs, as indicated for the ratio of [IAA^-^]_cyt_/[IAA^-^]_apo_ = 1 by the red line in **Figure 5b**. The different settings hardly affected the membrane voltage and E_K_. Both varied by ∼ 0.4 mV between the extremes (**Figure 5c,d**). In terms of [K^+^]_cyt_, this would be a variation of 1-2% (range 100-102 mM). In contrast, E_A_ varied by ∼ 4.5 mV, which was equivalent to a ∼ 20% difference in [A^-^]_cyt_ between low-*g*_*PIN*_/high-*g*_*ABC*_ and high-*g*_*PIN*_/low-*g*_*ABC*_ (**Figure 5e**). The low-*g*_*PIN*_/high-*g*_*ABC*_ condition consumed 3-4% more ATP than the high-*g*_*PIN*_/low-*g*_*ABC*_ condition (**Figure 5f**). This difference might be considered as small, but especially in metabolically highly active tissues (*e*.*g*., during growth), ATP may become a limiting factor and transmembrane transport processes must be supported by other energy sources (van Dongen *et al*., 2003; Gajdanowicz *et al*., 2011; Dreyer *et al*., 2017).

**Figure 5.**
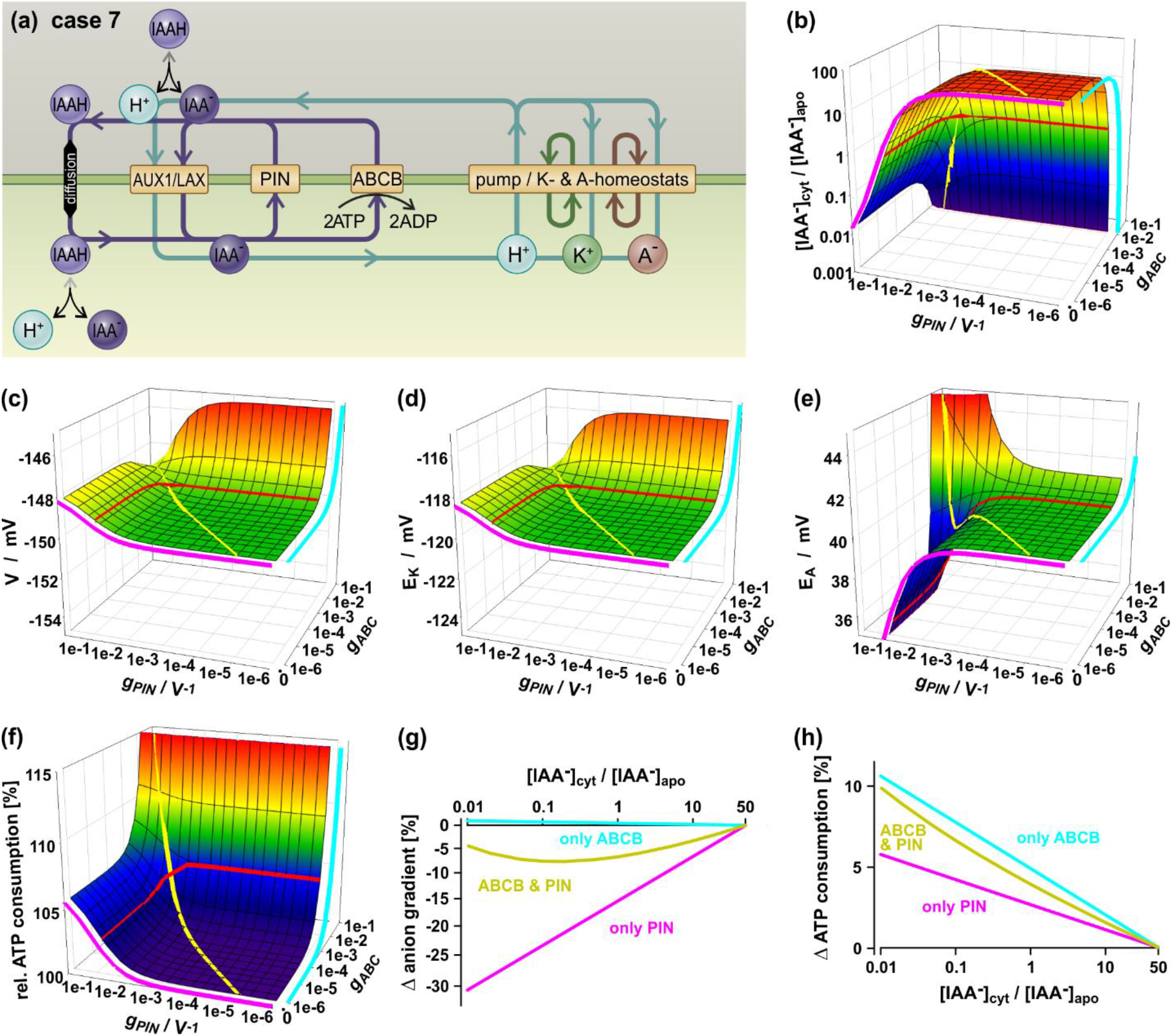
Coupling the activities of PINs and ABC transporters may mitigate the adverse effects of these transporter types on anion homeostasis and energy consumption. (a) Case 7: Schematic representation of the transporter network including ABC, PIN and AUX1 in addition to the natural IAAH diffusion and the K and A homeostats and the proton pump. The colored arrows and cycles indicate the net fluxes via the different transport pathways in steady state. (b-h) Steady state conditions of case 7 with *g*_*DIF*_+*g*_*AUX1*_ = 0.05 V^-1^. The curves in magenta show the values in steady state for scenarios in the absence of ABC-transporters (only PIN) while the curves in cyan indicate those in the absence of PINs (only ABC). The yellow curves specify a scenario, in which the activities of PINs and ABC-transporters are coupled with a constant ratio. (b-e) Dependency of the auxin gradient [IAA^-^]_cyt_/[IAA^-^]_apo_ (b), the membrane voltage *V* (c), the potassium gradient *E*_*K*_ (d), and the anion gradient *E*_*A*_ (e) on the activity of PIN and ABC. (f) Relative ATP consumption by the proton pump and the ABC transporter as a function of PIN and ABC activity. For reference, the ATP consumption of the network without auxin transporters (only K and A homeostats with pump) was set to 100%. The red curves in (b-f) show the values for the different (*g*_*PIN*_|*g*_*ABC*_)-pairs leading to [IAA^-^]_cyt_/[IAA^-^]_apo_ = 1. (g,h) Relative change of the transmembrane anion gradient (g) and the consumption of ATP (h) when a defined auxin gradient ([IAA^-^]_cyt_/[IAA^-^]_apo_) is adjusted.

The trade-off between (i) rapid adjustment of the desired auxin-gradient, (ii) lower ATP consumption for homeostasis (advantage PIN, disadvantage ABCB in both), and (iii) less interference with general anion homeostasis (disadvantage PIN, advantage ABCB) could be optimized by ABCB-PIN coupling that would synchronize the activity of both transporters. For illustration, we simulated that the activities of PIN and ABCB changed equally by the same factor (**Figure 5b-f**, yellow lines). In fact, this helped to mitigate the side effects. When we considered the detrimental change in anion gradient as a function of the desired [IAA^-^]_cyt_/[IAA^-^]_apo_ ratio, we found that the combination of ABCB and PIN performed better than PIN alone (**Figure 5g**). Similarly, the ABCB/PIN combination performed better than ABCB alone in terms of ATP consumption (**Figure 5h**). Thus, plants very likely employ two different transporter types for IAA^-^ export, PIN and ABCB, to mitigate the drawbacks of the individual transporters. Coupling between both could facilitate the fine-tuning of their activities required for the homeostatic control of auxin.

## Discussion

The ultimate goal of this work was to explore the possibility that an auxin homeostat functions in the establishment of apoplastic auxin gradients. Further, we aimed to understand the contribution and relevance of different auxin transporters to such an auxin homeostat and to test the option that individual members of auxin exporter classes act in a coordinated manner. In order to do so, we adopted a stepwise, unicellular, computational approach that was recently established for K^+^, A^-^, and Ca^2+^ homeostasis (Dreyer, 2021; Dindas *et al*., 2021; Dreyer *et al*., 2022; Li *et al*., 2024). Our study therefore contributes to the interdisciplinary quantitative plant biology (Morris & Ten Tusscher, 2021; Autran *et al*., 2021) approach to understand the multiple roles of auxin in plants. Not surprisingly, our simulations clearly indicated that neither auxin diffusion alone (case 1) nor diffusion in combination with AUX1/LAX (case 2) was sufficient to effectively control the apoplastic auxin concentration (**Figure 1**). Interestingly, addition of an IAA^-^ exporter, either an ABCB (case 3; **Figure 2**) or a PIN (cases 5 and 6; **Figure 3**), allowed for an adjustment of apoplastic auxin gradients by tuning the activity of the involved transporters (AUX1/LAX and ABCB/PINs). Importantly, unlike an ABCB, PINs could not increase the auxin gradient beyond the diffusion-determined limit (cyan line in **Figure 3b**). In summary, we conclude that an efficient auxin homeostat requires the existence of a combination of two different transport processes, diffusion and active transport, thus allowing the cell to gain control over the transmembrane auxin gradient.

Nevertheless, this advantage came for both tested exporter classes with a cost: like in any homeostat, also for ABCBs and PINs, this was an energy-consuming “auxin cycling” across the membrane (**Figures 2** and **3**, left panels) as already shown for K^+^ and anion homeostats (Dreyer, 2021). Further, the activity of ABCBs affected other homeostats in the membrane by either increasing or decreasing the *pmf*, respectively, that had obviously also an effect on K^+^- and A^-^-homeostasis (**Figure 2**; **Figure S2**). Like ABCBs, also PINs converted the static scenario of case 2 (diffusion plus AUX1/LAX) into a dynamic one (**Figure 3**; **Figures S4-S6**). In analogy to ABCBs, activity of PIN depolarized the membrane, which in turn affected K^+^ and anion homeostasis. Interestingly, our analyses indicated that for the more likely scenario of less selectively PINs (permeable beside IAA-for other anions; Deslauriers & Spalding, 2021), PIN activity interfered much stronger with general anion homeostasis than was the case with an IAA^-^-selective uniporter or an IAA^-^-specific ABCB (**Figure 4**).

A hitherto puzzling question was the relevance of the co-existence of two independent auxin-exporting systems, for which *independent* and *interactive* modes of action had been suggested (Bandyopadhyay *et al*., 2007; Blakeslee *et al*., 2007; Spalding, 2013; Geisler *et al*., 2017; Geisler, 2021; Deslauriers & Spalding, 2021; Mellor *et al*., 2022). A combined scenario with AUX1/LAX, ABCB, and PIN has now provided new insights that result directly from the thermodynamic properties of the system (case 7; **Figure 5**). As in the cases before, the transporter network enabled the proper adjustment of the transmembrane auxin gradient by regulating the transporter activities, at the expense of an energy-consuming auxin cycling across the membrane, which also affected K^+^ and anion homeostasis. However, these detrimental effects were not always of the same magnitude. Importantly, the usage of two different transporter types for IAA^-^ export, PIN and ABCB, mitigated the drawbacks of the individual transporters (ABCB: specificity > speed; PIN: speed > specificity) (**Figure 5**).

An unexpected finding was that ABCB-PIN coupling (i.e. synchronizing the activity of both transporters) apparently allows to optimize the trade-offs between speed of auxin gradient adjustment on one hand and ATP consumption and interference with general anion homeostasis on the other. Such a coupling of ABCBs and PINs — that *per se* in an *independent* action (Geisler *et al*., 2017) are sufficient for auxin homeostasis (Figures 2-3) — would fall in the category of a *cooperative interactive* action (Spalding, 2013; Geisler *et al*., 2017) and is supported theoretically (Spalding, 2013; Geisler *et al*., 2017; Mellor *et al*., 2022) and experimentally (Blakeslee *et al*., 2007; Deslauriers & Spalding, 2021). Overlapping expression profiles (Mellor *et al*., 2022) and successful co-immunoprecipitation (Bandyopadhyay *et al*., 2007; Blakeslee *et al*., 2007) suggest that such a functional coupling may be caused by physical ABCB-PIN interaction in which one of the auxin transporters acts as a regulator of the other, which functions primarily as a transport catalyst. Yet, neither modelling (Mellor *et al*., 2022) nor biochemical data (Blakeslee *et al*., 2007; Deslauriers & Spalding, 2021) have allowed for a dissection which transporter class functions as regulator or catalyst, respectively. However, for Arabidopsis ABCB19 such a regulatory role is supported by the finding that it was found to stabilize PIN1 protein membrane stability (Noh *et al*., 2003). Mammalian ABCs are well-known regulators of secondary active transport systems, including channels (Spalding, 2013; Aryal *et al*., 2015; Geisler *et al*., 2017). A prominent example is the sulphonylurea receptor (SUR/ ABCC8;) that associates with and regulates potassium channels, Kir6.1 or Kir6.2 (Principalli *et al*., 2015).

Interestingly, *cooperative, interactive* ABCB-PIN coupling was found to also enhance substrate (IAA^-^) specificity (Blakeslee *et al*., 2007; Spalding, 2013). Such an event might provide a rational for the optimization between speed of auxin gradient adjustment and ATP consumption and interference with general anion homeostasis found for ABCB-PIN coupling. Beside the here sketched scenario of *cooperative* ABCB-PIN functionality by protein-protein interaction, such a coupling of auxin transport between ABCBs and PINs could also be caused by a common regulatory module, such as phosphorylation by overlapping protein kinases. Transport of both ABCBs and PINs was demonstrated to be likewise regulated by overlapping subsets of AGC class XIII kinases, including PINOID and Phot1 (Christie *et al*., 2011; Henrichs *et al*., 2012).

Another remarkable outcome of our modelling approach was that the apparent relevance of each transporter class for auxin homeostasis roughly recapitulates the current picture of auxin transporter evolution (Vosolsobě *et al*., 2020; Geisler, 2021; Carrillo-Carrasco *et al*., 2023). In that, ABCB (and here not further discussed, ER-based PIN-LIKES, PILS; Feraru *et al*., 2012) transporters are ancient auxin transporters and thus already exist in chlorophytes, while PINs — although they seem to appear in some chlorophytes (Skokan *et al*., 2019; Carrillo-Carrasco *et al*., 2023) — can be safely associated only with charophytic algae (Carrillo-Carrasco *et al*., 2023). The origin of AUX1/LAXs, showing a more fragmentary distribution over charophytes and chlorophytes is less clear (Vosolsobě *et al*., 2020; Geisler, 2021; Carrillo-Carrasco *et al*., 2023).

In agreement with the fact that diffusion alone does not allow for control of auxin homeostasis (case 1), we are not aware of any organism that does not contain a minimum of one transporter class. Diffusion plus ABCB, for example, allows the control of auxin gradients across the PM, but their control is limited (case 3). In particular, the slow diffusion process hinders the rapid establishment of desired auxin gradients. In that respect it is worth mentioning that the early divergent charophyte algae *Chlorokybus atmophyticus* has ABCBs but apparently no PIN or AUX1/LAX (Vosolsobě *et al*., 2020; Geisler, 2021; Carrillo-Carrasco *et al*., 2023). The addition of AUX1/LAX was thus a major evolutionary event providing good homeostatic control overcoming the diffusion limit (case 2). This is underlined by a recent experimental quantification of auxin fluxes using a unicellular (protoplast)-based system, estimating that ca. 75% of the PM auxin influx is provided by AUX1, while 20% was assigned to other importers and only 5% is due to diffusion (Rutschow *et al*., 2014). Still, it is worth mentioning that in our study we assumed that the symporter AUX1/LAX is electroneutral and transports 1 H^+^ per 1 IAA^-^. In this case, and in the absence of an <u>imp</u>orting ABCB transporter, E_IAA_ was never more positive than E_H_, which means that the auxin gradient [IAA^-^]_cyt_/[IAA^-^]_apo_ was never larger than [H^+^]_apo_/[H^+^]_cyt_, *i*.*e*. ∼50 in physiological conditions. If, on the other hand, AUX1/LAX were electrogenic (n H^+^ / 1 IAA^-^, n > 1), the upper limit of the auxin gradient could be greater, but this is unlikely to be physiologically relevant. Moreover, maintaining IAA, H^+^, K^+^, and A^-^-cycles would be more energy-intensive in this case. Under homeostatic steady state conditions, for every IAA^-^ transported by AUX1/LAX, n protons are co-transported, which have to be pumped out of the cell again, consuming ATP. Considering the effects of this energy consumption on the other homeostats, the most efficient transport process would be for n = 1. The same conclusion has previously been worked out for putative electroneutral and electrogenic H^+^/K^+^ antiporters (Dreyer, 2021). Thus, the potential advantage of an electrogenic AUX1/LAX to be able to import auxin beyond the diffusion limit most likely does not outweigh the disadvantage of higher permanent energy consumption.

As ABCB transporters are faster than diffusion but significantly slower than H^+^/IAA^-^ symport, this scenario creates an obvious limit in the export of IAA^-^, which was filled with the next evolutionary event, the addition of PINs (cases 5 and 6). The interplay between diffusion-AUX1/LAX-ABCB and PIN enabled an excellent homeostatic control. Still, a valid question that arises is why (ancient) ABCBs have not yet been eliminated? Our data now indicate that ABCB-PIN pairs have advantages that PIN alone does not have. Thus, one might imagine that one day ABCBs may become obsolete if evolution creates an IAA^-^-selective PIN that does not allow the passage of smaller anions. However, from a biophysical point of view, considering the mode of operation of selectivity filters in ion channels (Doyle *et al*., 1998; Gouaux & MacKinnon, 2005), this may not be achievable for a large anion such as Indole-3-acetic acid.

In summary, our simulations of transporter networks show that individual auxin exporters (ABCB or PIN) are fully capable of creating proper auxin gradients, however, that functional ABCB-PIN coupling facilitates the fine-tuning of their activities required for a homeostatic control of auxin. Overall, our findings are in agreement with experimental studies (Blakeslee *et al*., 2007) but also a recent modeling study (Mellor *et al*., 2022) that focused on the generation of auxin fluxes rather than on the creation of gradients as performed here. Because the establishment of transmembrane auxin gradients is the basis of auxin fluxes, future modeling studies might now combine both complementary approaches and aim at integrating transporter polarity and adjusting transport capacities. Results from modeling approaches provide well-founded hypotheses for wet lab transport studies using functional reconstitution of transporters allowing for a proper dissection of transporter coupling. Our findings on the dynamics of apoplastic auxin distribution contribute to an understanding of the complexities of plant growth and development. This knowledge might be an essential basis for applications in various agricultural contexts, including crop improvement and the development of strategies for optimizing plant architecture and productivity.

## Acknowledgments

This work was supported by grants from the Swiss National Funds (project 31003A_165877 and 310030_197563 to MG) and the Agencia Nacional de Investigación y Desarrollo de Chile (ANID; grants Fondecyt Regular No. 1220504, and Anillo No. ATE220043 to ID).

## Author contributions

MG developed the idea of the project. MG and ID conceptionally designed the project. ID performed the mathematical modeling and the computational simulations. MG and ID prepared the figures, wrote the manuscript and approved the final version.

## Data availability

Further details about the expression and function of auxin transporters is available from Markus Geisler and further details about the modeling approach from Ingo Dreyer.

## Supporting Information

Additional Supporting Information may be found online in the Supporting Information section at the end of the article.

**Fig. S1** Voltage-dependence of IAA^-^ transporting ABCB transporters.

**Fig. S2** IAA^-^ importing ABCB-transporters can increase the transmembrane auxin gradient (case 4).

**Fig. S3** Effect of adjusting the activities of the ABCB-transport and the AUX1/LAX-transporter on membrane voltage, and equilibrium voltages for K^+^ and anions in cases 3 and 4.

**Fig. S4** Results for the scenario shown in Figure 3 in cases of strictly IAA^-^-selective PIN transporters.

**Fig. S5** Results for the scenario shown in Figure 3 in cases of non-selective PIN transporters.

**Fig. S6** Impact of the PIN-selectivity on cellular homeostasis.

**Fig. S7** Selectivity of PIN does not affect membrane voltage and K^+^ homeostasis.

**Methods S1** Mathematical representation of the steady state conditions of the systems considered in this study.

