## Supplemental Figures and Methods for "An auxin homeostat allows plant cells to establish and control defined transmembrane auxin gradients"

**Fig. S1**

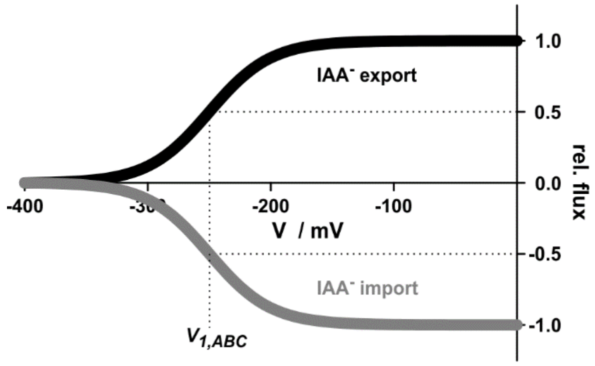


**Fig. S1** Voltage-dependence of $IAA^-$ transporting ABCB transporters. In the absence of substantial electrophysiological data, the ABCB transporters were modelled analogous to the plasma membrane $H^+$-ATPase (for details see Material and Methods). Two scenarios were considered: (i) The ABCB-transporter mediates an $IAA^-$ import (grey, negative flux) or (ii) an $IAA^-$ export (black, positive flux).

**Fig. S2**

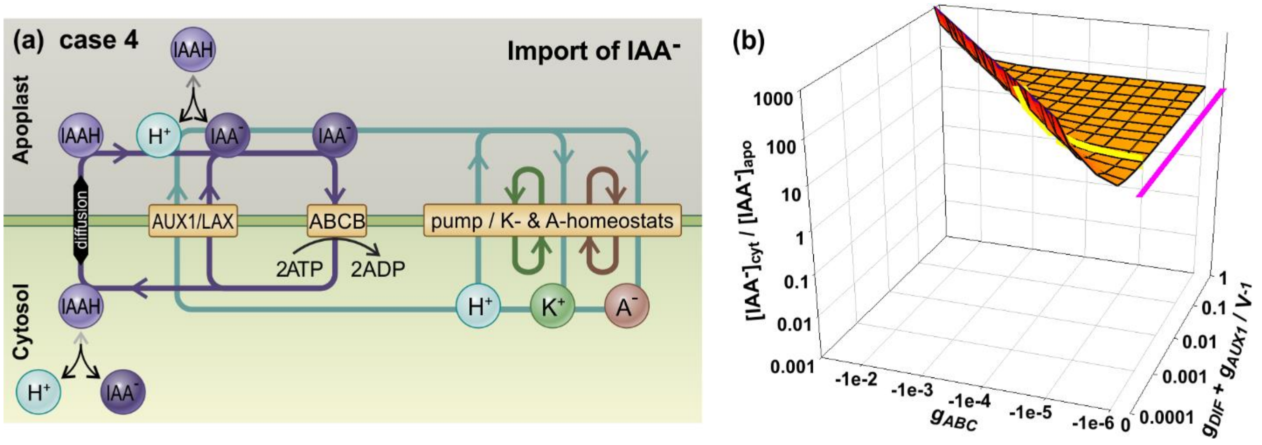


**Fig. S2** $IAA^-$ importing ABCB-transporters can increase the transmembrane auxin gradient (case 4). (a) Schematic representation of the transmembrane transport processes. ABCB-transporters import $IAA^-$, which is compensated in steady state by IAAH/$H^+$-$IAA^-$ efflux by diffusion or via AUX1/LAX. The involved proton fluxes in this cycling feeds back on the $H^+$-pump dependent homeostats for potassium ($K^+$) and anions ($A^-$). Substrate cycling is indicated by the colored arrows. (b) Transmembrane $[IAA^-]_{cyt}/[IAA^-]_{apo}$ gradient in steady state. The magenta line shows the state in the absence of ABCB transporter activity, which is determined by diffusion of IAAH and pH-dependent (de)protonation reactions. The yellow line depicts values for which $g_{DIF}+g_{AUX1} = 10^{-3}\ V^{-1}$ to illustrate exemplarily a lower limit determined by transporter-independent IAAH-diffusion across the membrane ($g_{AUX} = 0\ V^{-1}$).

#### Fig. S3

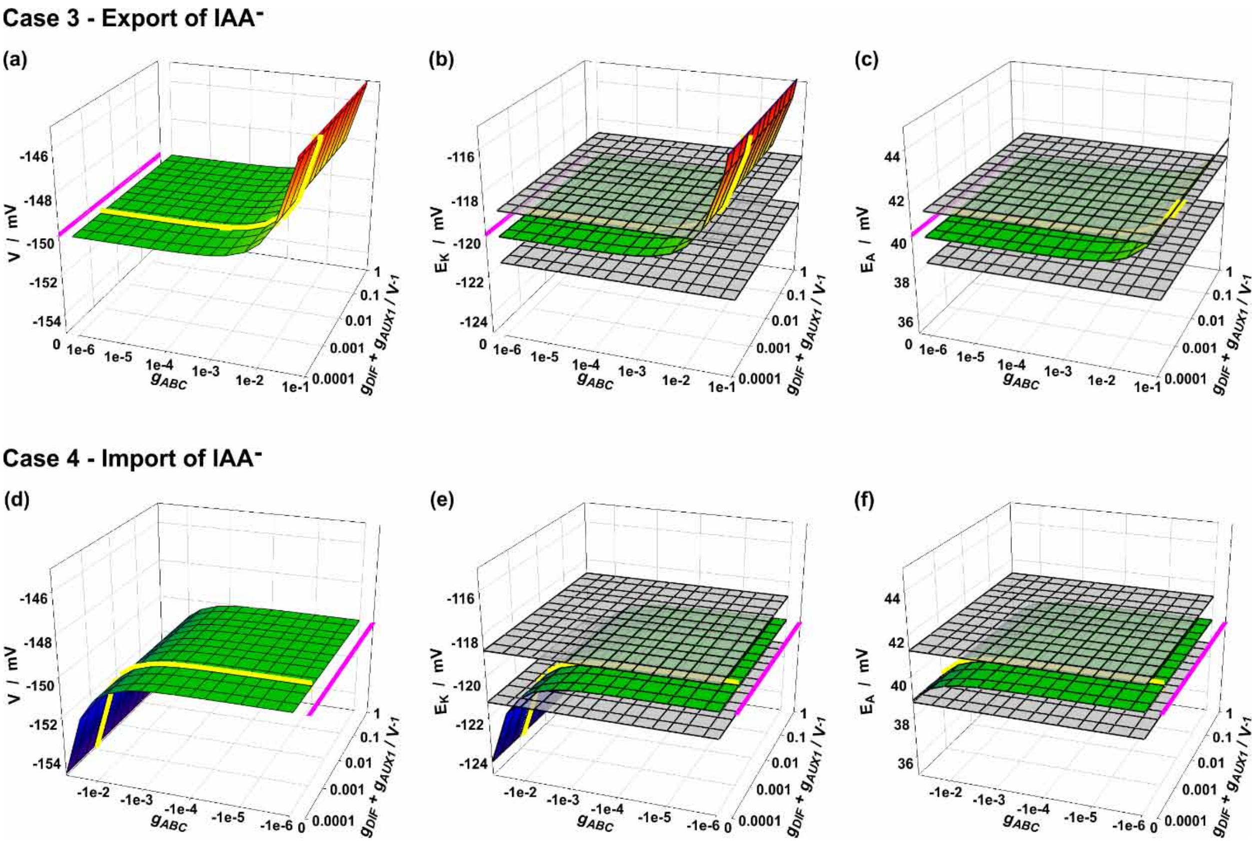


**Fig. S3** Effect of adjusting the activities of the ABCB-transport and the AUX1/LAX-transporter on membrane voltage, $V$ (a, d), and equilibrium voltages for $K^+$, $E_K$ (b, e), and anions, $E_A$ (c, f) in cases 3 (a-c) and 4 (d-f). The cyan lines show the conditions in the absence of ABCB transporter activity, which is determined by diffusion of IAAH and pH-dependent (de)protonation reactions. The yellow lines depict values for which $g_{DIF}$+$g_{AUX1}$ = $10^{-3}$ $V^{-1}$ to illustrate a lower limit determined by transporter-independent IAAH-diffusion across the membrane ($g_{AUX}$ = 0 $V^{-1}$). The grey surfaces indicate the tolerance limits for $E_K$ and $E_A$ that were set to $|\Delta E_K|$ = 1.25 mV and $|\Delta E_A|$ = 1.25 mV.

**Fig. S4**

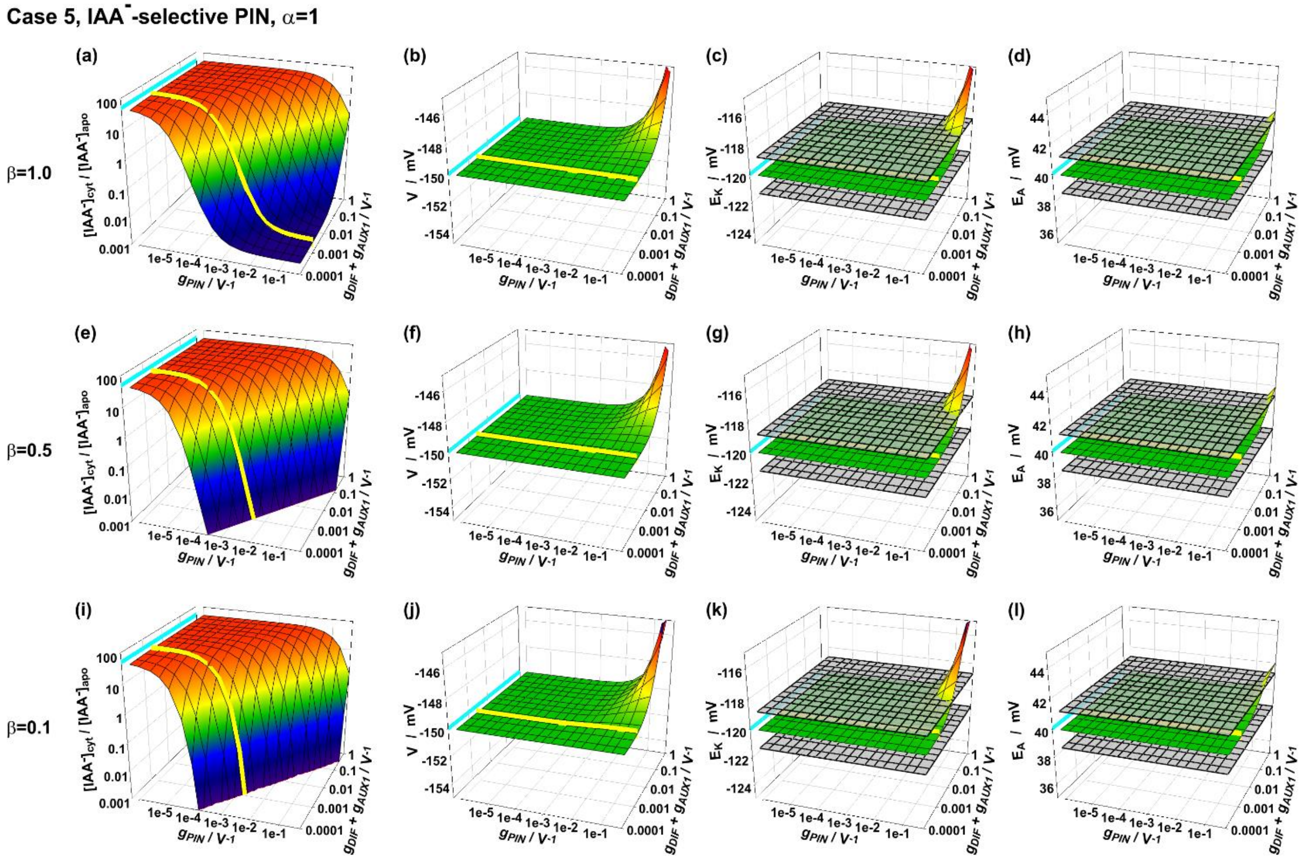


**Fig. S4** Results for the scenario shown in Figure 3 in cases of strictly $IAA^-$-selective PIN transporters (case 5, $\alpha$ = 1). The parameter $\beta$ was varied between 0.1 and 1: $\beta$ = 1.0, the driving force for $IAA^-$-transport came exclusively from the auxin gradient (a-d), $\beta$ = 0.5, $IAA^-$-transport was driven by the auxin and the anion ($A^-$) gradient (e-h), and $\beta$ = 0.1, $IAA^-$-transport was slightly driven by the auxin gradient but mostly by the gradient of the dominant anion $A^-$, e.g. $Cl^-$ or $NO_3^-$ (i-l). Shown are the transmembrane $[IAA^-]_{cyt}/[IAA^-]_{apo}$ gradients (a, e, i), membrane voltage, *V* (b, f, j), and equilibrium voltages for $K^+$, $E_K$ (c, g, k), and anions, $E_A$ (d, h, l) as a function of the PIN and AUX1/LAX/diffusion transporter activities. The cyan lines show the steady state in the absence of PIN transporter activity, which is determined by diffusion of IAAH and pH-dependent (de)protonation reactions. The yellow lines depict values for which $g_{DIF}+g_{AUX1}$ = $10^{-3}$ $V^{-1}$ to illustrate exemplarily a lower limit determined by transporter-independent IAAH-diffusion across the membrane ($g_{AUX}$ = 0 $V^{-1}$).

#### Fig. S5

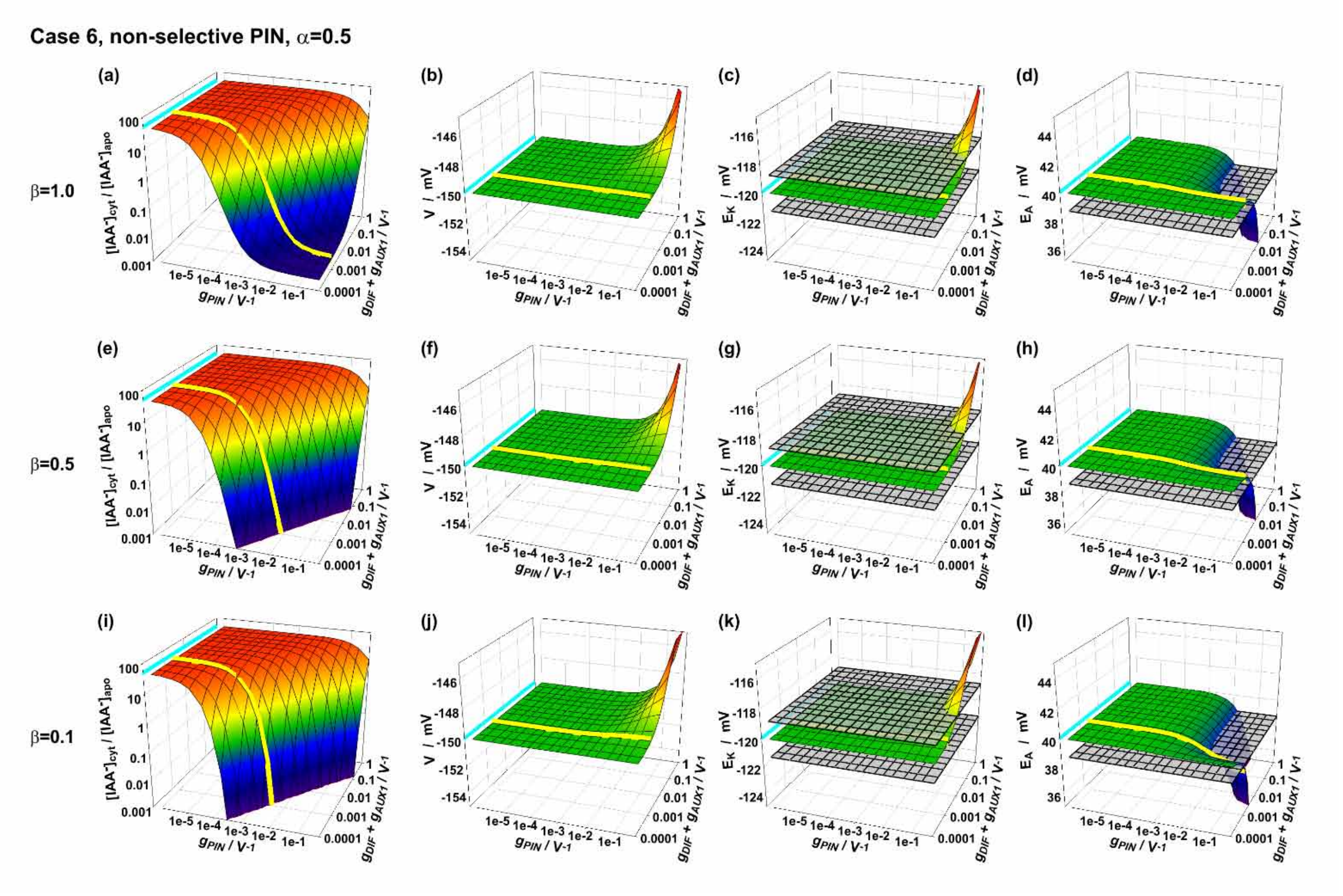


**Fig. S5** Results for the scenario shown in Figure 3 in cases of non-selective PIN transporters (case 6, exemplarily $\alpha = 0.5$). The parameter $\beta$ was varied between 0.1 and 1: $\beta = 1.0$, the driving force for $IAA^-$-transport came exclusively from the auxin gradient (a-d), $\beta = 0.5$, $IAA^-$-transport was driven by the auxin and the anion ($A^-$) gradient (e-h), and $\beta = 0.1$, $IAA^-$-transport was slightly driven by the auxin gradient but mostly by the gradient of the dominant anion $A^-$, e.g. $Cl^-$ or $NO_3^-$ (i-l). Shown are the transmembrane $[IAA^-]_{cyt}/[IAA^-]_{apo}$ gradients (a, e, i), membrane voltage, *V* (b, f, j), and equilibrium voltages for $K^+$, $E_K$ (c, g, k), and anions, $E_A$ (d, h, l) as a function of the PIN and AUX1/LAX/diffusion transporter activities. The cyan lines show the steady state in the absence of PIN transporter activity, which is determined by diffusion of IAAH and pH-dependent (de)protonation reactions. The yellow lines depict values for which $g_{DIF}+g_{AUX1} = 10^{-3}\ V^{-1}$ to illustrate exemplarily a lower limit determined by transporter-independent IAAH-diffusion across the membrane ($g_{AUX} = 0\ V^{-1}$).

### Fig. S6

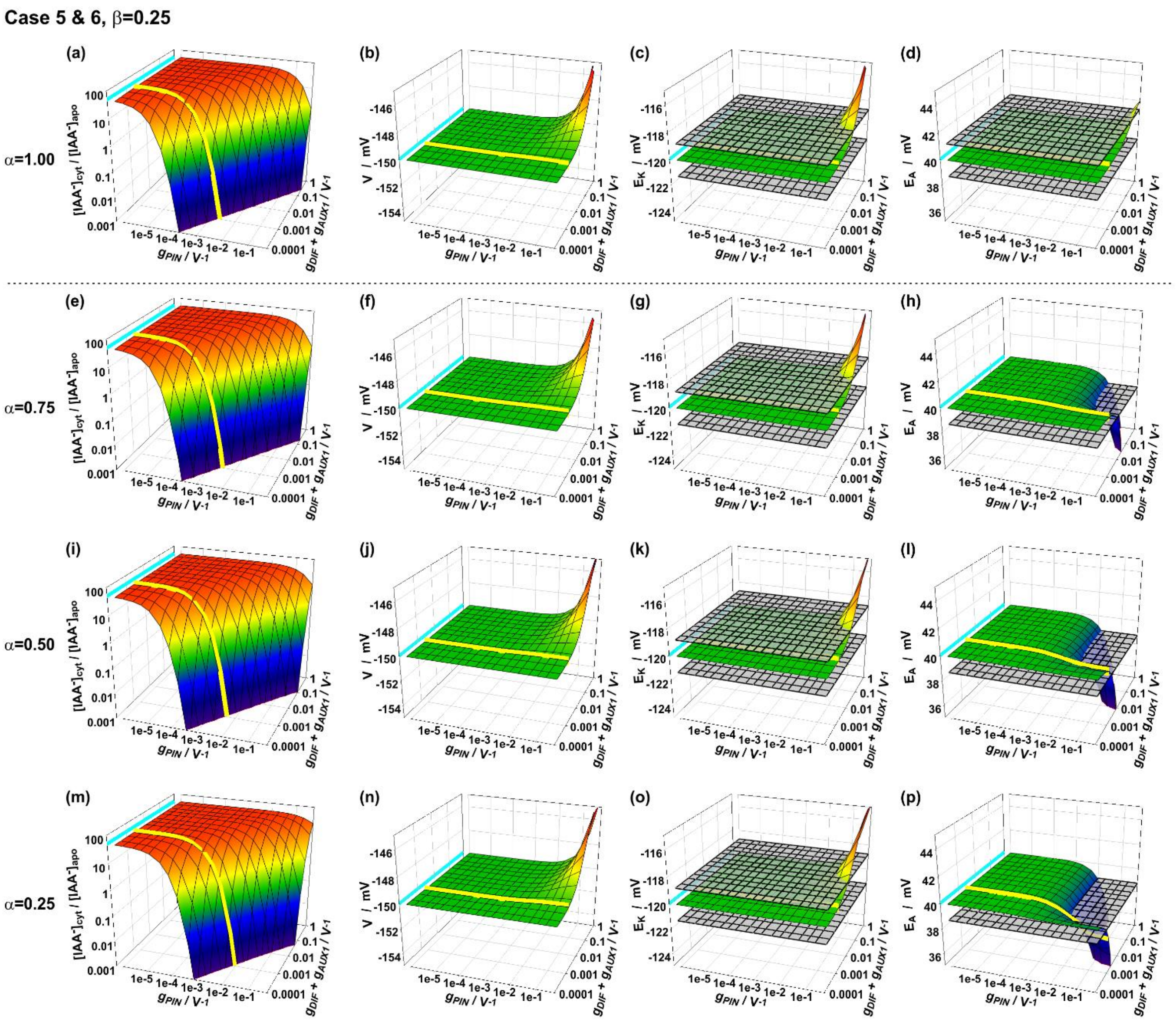


**Fig. S6** Impact of the PIN-selectivity on cellular homeostasis. Results for the scenario shown in Figure 3 for a fixed parameter $\beta$ (exemplarily $\beta$ = 0.5), while the parameter $\alpha$ was varied: (a-d) $\alpha$ = 1.00, PIN is 100% (exclusively) permeable for $IAA^-$, and 0% (impermeable) for the dominant small anion $A^-$, e.g. $Cl^-$ or $NO_3^-$. (e-h) $\alpha$ = 0.75, PIN is largely permeable for $IAA^-$, but also slightly for $A^-$. (i-l) $\alpha$ = 0.50, PIN is just as permeable for $IAA^-$ as for $A^-$. (m-p) $\alpha$ = 0.25, PIN is better permeable for $A^-$ than for $IAA^-$. Shown are the transmembrane $[IAA^-]_{cyt}/[IAA^-]_{apo}$ gradients (a, e, i, m), membrane voltage, $V$ (b, f, j, n), and equilibrium voltages for $K^+$, $E_K$ (c, g, k, o), and anions, $E_A$ (d, h, l, p) as a function of the PIN and AUX1/LAX/diffusion transporter activities. The cyan lines show the steady state in the absence of PIN transporter activity, which is determined by diffusion of IAAH and pH-dependent (de)protonation reactions. The yellow lines depict values for which $g_{DIF}+g_{AUX1} = 10^{-3}\ V^{-1}$ to illustrate exemplarily a lower limit determined by transporter-independent IAAH-diffusion across the membrane ($g_{AUX} = 0\ V^{-1}$).

**Fig. S7**

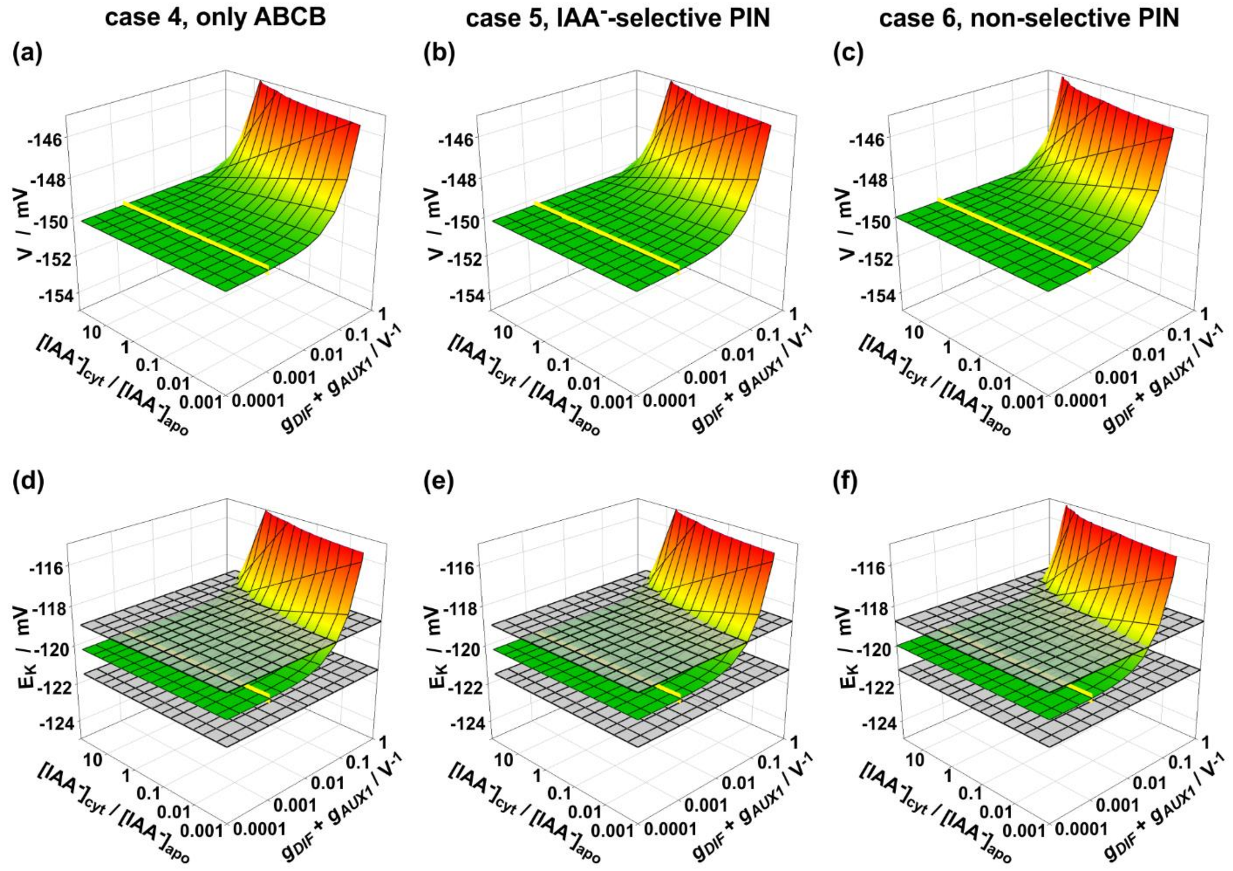


**Fig. S7** Selectivity of PIN does not affect membrane voltage and $K^+$ homeostasis. Comparison of the steady values of the membrane voltage V (a-c) and $E_K$ $[= \frac{RT}{F} \cdot \ln\left(\frac{[K^+]_{apo}}{[K^+]_{cyt}}\right)]$ (d-f) depending on auxin diffusion ($g_{DIF}$+$g_{AUX1}$) and the desired transmembrane auxin gradient ($[IAA^-]_{cyt}/[IAA^-]_{apo}$) in case 4 with $IAA^-$-exporting ABCB transporters (a,d), case 5 ($\alpha$ = 1, $\beta$ = 1) with $IAA^-$-selective PINs (b,e), and case 6 ($\alpha$ = 0.5, $\beta$ = 1) with non-selective PINs (c,f). The grey surfaces indicate thresholds at which the transmembrane anion gradient changes by 5% ($|\Delta E_K|$ = 1.25 mV) compared to the reference condition without auxin transporters. The yellow lines indicate values for which $g_{DIF}+g_{AUX1} = 10^{-3}\ V^{-1}$ to illustrate exemplarily a lower limit determined by transporter-independent IAAH-diffusion across the membrane ($g_{AUX} = 0\ V^{-1}$).

**Methods S1** Mathematical representation of the steady state conditions of the systems considered in this study.

$$\begin{cases} 0 = i_P(V) + g_{KC} \cdot (V - E_K) + 2 \cdot g_{KHs} \cdot (2 \cdot V - E_H - E_K) + g_{AC} \cdot (V - E_A) + g_{HA} \cdot (V - 2 \cdot E_H + E_A) + g_{PIN} \cdot (V - \beta \cdot E_{IAA} - (1-\beta) \cdot E_A) - g_{ABC} \cdot i_{ABC}(V) \\ 0 = g_{KC} \cdot (V - E_K) + g_{KHs} \cdot (2 \cdot V - E_H - E_K) - g_{KHa} \cdot (E_K - E_H) \\ 0 = -g_{AC} \cdot (V - E_A) + g_{HA} \cdot (V - 2 \cdot E_H + E_A) - (1-\alpha) \cdot g_{PIN} \cdot (V - \beta \cdot E_{IAA} - (1-\beta) \cdot E_A) \\ 0 = g_{DIF} \cdot (E_{IAA} - E_H) + g_{AUX} \cdot (E_{IAA} - E_H) - \alpha \cdot g_{PIN} \cdot (V - \beta \cdot E_{IAA} - (1-\beta) \cdot E_A) + g_{ABC} \cdot i_{ABC}(V) \end{cases} \tag{S1}$$

arranged in vector and matrix form:

$$M \cdot \begin{pmatrix} V \\ E_K \\ E_A \\ E_{IAA} \end{pmatrix} = \vec{A}$$

with:

$$M = \begin{pmatrix} g_{KC} + 4 \cdot g_{KHs} + g_{AC} + g_{HA} + g_{PIN} & -g_{KC} - 2 \cdot g_{KHs} & g_{HA} - g_{AC} - (1-\beta) \cdot g_{PIN} & -\beta \cdot g_{PIN} \\ g_{KC} + 2 \cdot g_{KHs} & -g_{KC} - g_{KHs} - g_{KHa} & 0 & 0 \\ g_{HA} - g_{AC} - (1-\alpha) \cdot g_{PIN} & 0 & g_{AC} + g_{HA} + (1-\alpha) \cdot (1-\beta) \cdot g_{PIN} & (1-\alpha) \cdot \beta \cdot g_{PIN} \\ -\alpha \cdot g_{PIN} & 0 & \alpha \cdot (1-\beta) \cdot g_{PIN} & g_{DIF} + g_{AUX} + \alpha \cdot \beta \cdot g_{PIN} \end{pmatrix}$$

and:

$$\vec{A} = \begin{pmatrix} -i_p + (2 \cdot g_{KHs} + 2 \cdot g_{HA}) \cdot E_H + g_{ABC} \cdot i_{ABC} \\ (g_{KHs} - g_{KHa}) \cdot E_H \\ 2 \cdot g_{HA} \cdot E_H \\ (g_{DIF} + g_{AUX}) \cdot E_H - g_{ABC} \cdot i_{ABC} \end{pmatrix}$$

The solution of this equation system is:

$$V = E_H - \frac{\frac{(2 \cdot g_{HA} + (g_{AC} - g_{HA}) \cdot \alpha) \cdot \beta \cdot g_{PIN} \cdot g_{ABC} \cdot i_{ABC}}{(g_{AC}+g_{HA}) \cdot (g_{DIF}+g_{AUX}+\alpha \cdot \beta \cdot g_{PIN}) + (1-\alpha) \cdot (1-\beta) \cdot (g_{DIF}+g_{AUX}) \cdot g_{PIN}} + i_p - g_{ABC} \cdot i_{ABC}}{\frac{4 \cdot g_{KHa} \cdot g_{KHs} + g_{KC} \cdot g_{KHa} + g_{KC} \cdot g_{KHs}}{g_{KC}+g_{KHa}+g_{KHs}} + \frac{(g_{AC}+g_{HA}) \cdot \alpha \cdot \beta \cdot (g_{DIF}+g_{AUX}) \cdot g_{PIN} + 4 \cdot g_{AC} \cdot g_{HA} \cdot (g_{DIF}+g_{AUX}+\alpha \cdot \beta \cdot g_{PIN}) + 2 \cdot g_{HA} \cdot (1-\alpha+1-\beta) \cdot (g_{DIF}+g_{AUX}) \cdot g_{PIN}}{(g_{AC}+g_{HA}) \cdot (g_{DIF}+g_{AUX}+\alpha \cdot \beta \cdot g_{PIN}) + (1-\alpha) \cdot (1-\beta) \cdot (g_{DIF}+g_{AUX}) \cdot g_{PIN}}} \tag{S2}$$

$$E_K = E_H + \frac{g_{KC} + 2 \cdot g_{KHs}}{g_{KC}+g_{KHa}+g_{KHs}} \cdot (V - E_H) \tag{S3}$$

$$E_A = E_H + (V - E_H) \cdot$$

$$\frac{\left((g_{AC}-g_{HA}) \cdot (g_{DIF}+g_{AUX}+\alpha \cdot \beta \cdot g_{PIN}) + (1-\alpha) \cdot (g_{DIF}+g_{AUX}) \cdot g_{PIN}\right) \cdot i_p - \left((g_{AC}-g_{HA}) \cdot (g_{DIF}+g_{AUX}) + \left(\left(2 \cdot g_{HA} + \frac{4 \cdot g_{KHa} \cdot g_{KHs} + g_{KC} \cdot g_{KHa} + g_{KC} \cdot g_{KHs}}{g_{KC}+g_{KHa}+g_{KHs}}\right) \cdot \beta + g_{DIF} + g_{AUX}\right) \cdot (1-\alpha) \cdot g_{PIN}\right) \cdot g_{ABC} \cdot i_{ABC}}{(2 \cdot g_{HA} + (g_{AC}-g_{HA}) \cdot \alpha) \cdot \beta \cdot g_{PIN} \cdot g_{ABC} \cdot i_{ABC} + (i_p - g_{ABC} \cdot i_{ABC}) \cdot \left((g_{AC}+g_{HA}) \cdot (g_{DIF}+g_{AUX}+\alpha \cdot \beta \cdot g_{PIN}) + (1-\alpha) \cdot (1-\beta) \cdot (g_{DIF}+g_{AUX}) \cdot g_{PIN}\right)}$$

(S4)

$$E_{IAA} = E_H + (V - E_H) \cdot \frac{\frac{4\cdot g_{KHa}\cdot g_{KHs}+g_{KC}\cdot g_{KHa}+g_{KC}\cdot g_{KHs}}{g_{KC}+g_{KHa}+g_{KHs}}\cdot\left((g_{AC}+g_{HA})+(1-\alpha)\cdot(1-\beta)\cdot g_{PIN}\right)\cdot g_{ABC}\cdot i_{ABC}+4\cdot g_{AC}\cdot g_{HA}\cdot g_{ABC}\cdot i_{ABC}+(g_{AC}-g_{HA})\cdot\alpha\cdot\beta\cdot g_{PIN}\cdot i_p+2\cdot g_{HA}\cdot g_{PIN}\cdot\left(\alpha\cdot i_p+(1-\alpha)\cdot(2-\beta)\cdot g_{ABC}\cdot i_{ABC}\right)}{\left(2\cdot g_{HA}+(g_{AC}-g_{HA})\cdot\alpha\right)\cdot\beta\cdot g_{PIN}\cdot g_{ABC}\cdot i_{ABC}+(i_p-g_{ABC}\cdot i_{ABC})\cdot\left((g_{AC}+g_{HA})\cdot(g_{DIF}+g_{AUX}+\alpha\cdot\beta\cdot g_{PIN})+(1-\alpha)\cdot(1-\beta)\cdot(g_{DIF}+g_{AUX})\cdot g_{PIN}\right)} \quad \text{(S5)}$$

Please note that $i_p$ and $i_{ABC}$ depend on the voltage. Thus, the solution of the equation system is an implicit solution. To get numerical values for $V$, $E_K$, $E_A$, and $E_{IAA}$ it still needs to be solved numerically for a given parameter set [$g_{KC}$, $g_{KHs}$, $g_{KHa}$, $g_{AC}$, $g_{HA}$, $g_{DIF}$, $g_{AUX}$, $g_{PIN}$, $g_{ABC}$, $E_H$].

#### Reference condition

As reference condition, we chose $g_{KC}$ = 1.000 $V^{-1}$, $g_{KHs}$ = 0.000V-1, $g_{KHa}$ = 0.138 $V^{-1}$, $g_{AC}$ = 0.687 $V^{-1}$, $g_{HA}$ = 0.427 $V^{-1}$, and $E_H$ = +98 mV. This resulted in the following steady-state values in the absence of auxin:

$$V - E_H + \frac{i_p(V)}{\frac{4\cdot g_{AC}\cdot g_{HA}}{g_{AC}+g_{HA}} + \frac{4\cdot g_{KHa}\cdot g_{KHs}+g_{KC}\cdot g_{KHa}+g_{KC}\cdot g_{KHs}}{g_{KC}+g_{KHa}+g_{KHs}}} = 0 = -150mV - 98mV + \frac{0.291}{1.174V^{-1}}$$

$$\Rightarrow V \approx -150mV$$

$$E_K = E_H + \frac{g_{KC}+2\cdot g_{KHs}}{g_{KC}+g_{KHa}+g_{KHs}}\cdot(V-E_H) = 98mV + 0.879\cdot(-150mV-98mV) \approx -120mV$$

$$E_A = E_H + \frac{g_{AC}-g_{HA}}{g_{AC}+g_{HA}}\cdot(V-E_H) = 98mV + 0.233\cdot(-150mV-98mV) \approx +40mV$$

#### Steady-state condition of the investigated cases

| | |
|---|---|
| Cases<br>1&2<br>DIF/AUX<br><br>$g_{ABC}$ = 0<br>$g_{PIN}$ = 0 | $V^{case\ 1\&2} = 98mV - \frac{i_p}{1.174V^{-1}} = -150mV$<br>$E_K^{case\ 1\&2} = 98mV + 0.879 \cdot (V - 98mV) = -120mV$<br>$E_A^{case\ 1\&2} = 98mV + 0.233 \cdot (V - 98mV) = +40mV$<br>$E_{IAA}^{case\ 1\&2} = 98mV$ |
| Cases<br>3&4<br>DIF/AUX<br>ABC<br><br>$g_{PIN}$ = 0 | $V^{case\ 3\&4} - 98mV + \frac{i_p - g_{ABC} \cdot i_{ABC}}{1.174V^{-1}} = 0$<br>$E_K^{case\ 3\&4} = 98mV + 0.879 \cdot (V - 98mV)$<br>$E_A^{case\ 3\&4} = 98mV + 0.233 \cdot (V - 98mV)$<br>$E_{IAA}^{case\ 3\&4} = 98mV - \frac{g_{ABC} \cdot i_{ABC}}{(g_{DIF} + g_{AUX})}$ |
| Cases<br>5&6<br>DIF/AUX<br>PIN<br><br>$g_{ABC}$ = 0 | $V^{case\ 5\&6} - 98mV + \frac{i_p}{0.121V^{-1} + \frac{1.114V^{-1} \cdot \alpha \cdot \beta \cdot (g_{DIF} + g_{AUX}) \cdot g_{PIN} + 1.173V^{-2} \cdot (g_{DIF} + g_{AUX} + \alpha \cdot \beta \cdot g_{PIN}) + 0.854V^{-1} \cdot (1 - \alpha + 1 - \beta) \cdot (g_{DIF} + g_{AUX}) \cdot g_{PIN}}{1.114V^{-1} \cdot (g_{DIF} + g_{AUX} + \alpha \cdot \beta \cdot g_{PIN}) + (1 - \alpha) \cdot (1 - \beta) \cdot (g_{DIF} + g_{AUX}) \cdot g_{PIN}}} = 0$<br>$E_K^{case\ 5\&6} = 98mV + 0.879 \cdot (V - 98mV)$<br>$E_A^{case\ 5\&6} = 98mV + \frac{0.260V^{-1} \cdot (g_{DIF} + g_{AUX} + \alpha \cdot \beta \cdot g_{PIN}) + (1 - \alpha) \cdot (g_{DIF} + g_{AUX}) \cdot g_{PIN}}{1.114V^{-1} \cdot (g_{DIF} + g_{AUX} + \alpha \cdot \beta \cdot g_{PIN}) + (1 - \alpha) \cdot (1 - \beta) \cdot (g_{DIF} + g_{AUX}) \cdot g_{PIN}} \cdot (V - 98mV)$<br>$E_{IAA}^{case\ 5\&6} = 98mV + \frac{0.260V^{-1} \cdot \beta + 0.854V^{-1}}{1.114V^{-1} \cdot (g_{DIF} + g_{AUX} + \alpha \cdot \beta \cdot g_{PIN}) + (1 - \alpha) \cdot (1 - \beta) \cdot (g_{DIF} + g_{AUX}) \cdot g_{PIN}} \cdot \alpha \cdot g_{PIN} \cdot (V - 98mV)$ |
| Case 7<br>DIF/AUX<br>PIN<br>ABC | $V^{case7} - 98mV + \frac{\frac{(0.854V^{-1} + 0.260V^{-1} \cdot \alpha) \cdot \beta \cdot g_{PIN} \cdot g_{ABC} \cdot i_{ABC}}{1.114V^{-1} \cdot (g_{DIF} + g_{AUX} + \alpha \cdot \beta \cdot g_{PIN}) + (1 - \alpha) \cdot (1 - \beta) \cdot (g_{DIF} + g_{AUX}) \cdot g_{PIN}} + i_p - g_{ABC} \cdot i_{ABC}}{0.121V^{-1} + \frac{1.114V^{-1} \cdot \alpha \cdot \beta \cdot (g_{DIF} + g_{AUX}) \cdot g_{PIN} + 1.173V^{-2} \cdot (g_{DIF} + g_{AUX} + \alpha \cdot \beta \cdot g_{PIN}) + 0.854V^{-1} \cdot (1 - \alpha + 1 - \beta) \cdot (g_{DIF} + g_{AUX}) \cdot g_{PIN}}{1.114V^{-1} \cdot (g_{DIF} + g_{AUX} + \alpha \cdot \beta \cdot g_{PIN}) + (1 - \alpha) \cdot (1 - \beta) \cdot (g_{DIF} + g_{AUX}) \cdot g_{PIN}}} = 0$<br>$E_K^{case\ 7} = E_H + 0.879 \cdot (V - E_H)$<br>$E_A^{c.7} = 98mV + \frac{(0.260V^{-1} \cdot (g_{DIF} + g_{AUX} + \alpha \cdot \beta \cdot g_{PIN}) + (1 - \alpha) \cdot (g_{DIF} + g_{AUX}) \cdot g_{PIN}) \cdot i_p - \left(0.260V^{-1} \cdot (g_{DIF} + g_{AUX}) + ((0.854V^{-1} + 0.121V^{-1}) \cdot \beta + g_{DIF} + g_{AUX}) \cdot (1 - \alpha) \cdot g_{PIN}\right) \cdot g_{ABC} \cdot i_{ABC}}{(0.854V^{-1} + 0.260V^{-1} \cdot \alpha) \cdot \beta \cdot g_{PIN} \cdot g_{ABC} \cdot i_{ABC} + (i_p - g_{ABC} \cdot i_{ABC}) \cdot (1.114V^{-1} \cdot (g_{DIF} + g_{AUX} + \alpha \cdot \beta \cdot g_{PIN}) + (1 - \alpha) \cdot (1 - \beta) \cdot (g_{DIF} + g_{AUX}) \cdot g_{PIN})} \cdot (V - 98mV)$<br>$E_{IAA}^{case7} = 98mV + \frac{0.121V^{-1} \cdot (1.114V^{-1} + (1 - \alpha) \cdot (1 - \beta) \cdot g_{PIN}) \cdot g_{ABC} \cdot i_{ABC} + 1.173V^{-2} \cdot g_{ABC} \cdot i_{ABC} + 0.260V^{-1} \cdot \alpha \cdot \beta \cdot g_{PIN} \cdot i_p + 0.854V^{-1} \cdot g_{PIN} \cdot (\alpha \cdot i_p + (1 - \alpha) \cdot (2 - \beta) \cdot g_{ABC} \cdot i_{ABC})}{(0.854V^{-1} + 0.260V^{-1} \cdot \alpha) \cdot \beta \cdot g_{PIN} \cdot g_{ABC} \cdot i_{ABC} + (i_p - g_{ABC} \cdot i_{ABC}) \cdot (1.114V^{-1} \cdot (g_{DIF} + g_{AUX} + \alpha \cdot \beta \cdot g_{PIN}) + (1 - \alpha) \cdot (1 - \beta) \cdot (g_{DIF} + g_{AUX}) \cdot g_{PIN})} \cdot (V - 98mV)$ |
